# Atypical spatial frequency dependence of visual metacognition among schizophrenia patients

**DOI:** 10.1101/853135

**Authors:** Ai Koizumi, Tomoki Hori, Brian Maniscalco, Makoto Hayase, Ryou Mishima, Takahiko Kawashima, Jun Miyata, Toshihiko Aso, Hakwan Lau, Hidehiko Takahashi, Kaoru Amano

## Abstract

Although altered early stages of visual processing have been reported among schizophrenia patients, how such atypical visual processing may affect higher-level cognition remains largely unknown. Here we tested the hypothesis that metacognitive performance may be atypically modulated by spatial frequency (SF) of visual stimuli among individuals with schizophrenia, given their altered magnocellular function. To study the effect of SF on metacognitive performance, we asked patients and controls to perform a visual detection task on gratings with different SFs and report confidence, and analyzed the data using the signal detection theoretic measure meta-d’. Control subjects showed better metacognitive performance after yes- (stimulus-presence) than after no- (stimulus-absence) responses (‘yes-response advantage’) for high SF (HSF) stimuli but not for low SF (LSF) stimuli. The patients, to the contrary, showed a ‘yes-response advantage’ not only for HSF but also for LSF stimuli, indicating atypical SF dependency of metacognition. An fMRI experiment using the same task revealed that the dorsolateral prefrontal cortex (DLPFC), known to be crucial for metacognition, shows activity mirroring the behavioral results: decoding accuracy of perceptual confidence in DLPFC was significantly higher for HSF than for LSF stimuli in controls, whereas this decoding accuracy was independent of SF in patients. While individuals without schizophrenia may flexibly adapt metacognitive computations across SF ranges, patients may employ a different mechanism that is independent of SF. Because visual stimuli of low SF have been linked to top-down processing in predictive coding, this may reflect atypical functioning in these processes in schizophrenia.

**Highlights:**

- Visual metacognition of controls was dependent on spatial frequency.
- Visual metacognition of schizophrenia patients was independent of spatial frequency.
- Patients and controls differently rely on the dorsolateral prefrontal cortex.
- Sensory inputs may reach metacognitive circuits in an atypical manner among patients.

## 1. Introduction

Schizophrenia is a neurodevelopmental psychopathology which is found in about 1% of the population (Regier et al., 1993; van Os and Kapur, 2009). A comprehensive understanding of how neural functions go awry with schizophrenia is still underway. Among schizophrenia patients, studies have demonstrated atypicality not only in their cognitive functions (Callicott et al., 2003; Goldman-Rakic and Selemon, 1997; Weinberger and Gallhofer, 1997) but also in their early sensory processing (Butler and Javitt, 2005; Butler et al., 2007; Javitt and Freedman, 2015; Saccuzzo et al., 1974). Although there has been speculation regarding the relationship between the two functions (Butler et al., 2007; Javitt and Freedman, 2015), how atypical sensory functions contribute to altered higher cognitive functions remains largely unknown. Here, we examine the potential interaction between early stage visual processing of spatial frequency (SF) and higher-order metacognitive function among schizophrenia patients.

One widely reported atypicality with schizophrenia in low-level visual function is the response profile across SF. SF is typically critical for gating sensory inputs for distinct visual computations. For example, while coarse visual inputs with lower SF rapidly frame the visual processing in a predictive top-down manner (Bar et al., 2006; Javitt and Freedman, 2015), fine visual inputs with higher SF contribute to more elaborated perceptual decisions e.g., object identification (Bar et al., 2006; Javitt and Freedman, 2015). Among schizophrenia patients, studies have shown reduced neural response to lower SF in visual cortices and some other areas, which is likely due to their magnocellular dysfunction (Butler and Javitt, 2005; Martinez et al., 2012; Martinez et al., 2008). Patients also showed hindered perceptual sensitivity to lower SF inputs in some (Butler and Javitt, 2005; Martinez et al., 2008; Slaghuis and Curran, 1999) but not all tasks (Keri et al., 2002).

Some studies have shown that at later stages of perceptual decision-making such as object recognition, patients put more weight on low SF input, while control subjects generally rely more on high SF input (Laprevote et al., 2010; Laprevote et al., 2013). One possibility is that, among schizophrenia patients, sensory inputs across SF ranges may get atypically wired to later stage visual processing, including metacognition.

We here examined whether low level visual properties of SF modulate higher-order metacognitive function in an atypical manner among schizophrenia patients. Metacognition involves the monitoring of one’s own cognition (Frith, 2012; Nelson and Narens, 1990). One critical metacognitive function is to evaluate the accuracy of one’s perceptual judgments (Fleming et al., 2012a; Lau and Rosenthal, 2011), which is carried out in higher cortical areas, especially the dorsolateral prefrontal cortex (DLPFC) (Cortese et al., 2016; Fleming et al., 2012b; Koizumi et al., 2016; Shekhar and Rahnev, 2018). This function enables one to discern between true and false percepts by creating an elevated sense of confidence in the former relative to the latter (Fleming et al., 2012a; Lau and Rosenthal, 2011).

Although recent studies have examined perceptual metacognitive performance among people with schizophrenia, without manipulation of SF, the results appear mixed and non-conclusive (Charles et al., 2017; Powers et al., 2017; Rouault et al., 2018). For example, patients showed lowered sensitivity for discriminating between correct versus incorrect percepts (i.e., metacognitive sensitivity, meta-d’) relative to controls (Charles et al., 2017) in conditions where patients also showed lowered perceptual sensitivity (d’). Thus, there remains a possibility that lowered metacognitive sensitivity was merely due to lowered perceptual sensitivity since perceptual sensitivity and metacognitive sensitivity generally covary (Fleming and Lau, 2014; Maniscalco and Lau, 2014). In line with this, Powers et al. (2017) reported that metacognitive sensitivity was unrelated with psychosis and/or hallucination when perceptual sensitivity (d’) was taken into account. Similarly, Rouault et al. (2018) showed that self-reported schizotypy had little contribution to metacognitive performance. One reason for these mixed results could be that schizophrenia patients are atypical not in their general metacognitive performance, but rather in how their metacognitive performance is modulated by features of the perceptual decision-making context, such as SF as investigated here.

We here examined metacognitive performance during a visual detection task with perceptual confidence rating using higher and lower SF stimuli among schizophrenia patients. During a detection task, metacognitive sensitivity for discriminating between correct versus incorrect percepts is typically better following stimulus-present responses (yes-responses) than for stimulus-absent responses (no-responses) among healthy individuals (Fleming and Dolan, 2010; Kanai et al., 2010) (see also our replication in Supplementary Figure 1). That is, it is generally easier to discriminate one’s correct versus incorrect yes-responses (i.e. by giving higher confidence ratings to hits vs false alarms) than to discriminate correct versus incorrect no-responses (i.e., correct rejections vs misses) when detecting a visual target.

An intuitive explanation for this phenomenon is that confidence is typically evaluated by reference to the amount of sensory evidence under consideration, and the amount of available sensory evidence differs for yes- and no-responses. In a detection task, a “yes” response indicates that the observer saw the target and therefore has access to sensory evidence for the target, and such sensory evidence can then be evaluated to determine confidence. By contrast, a “no” response indicates that the observer did not see the target and therefore likely has little-to-no sensory evidence pertaining to the target, which entails that the typical strategy of rating confidence by reference to the available sensory evidence is not tenable and an alternative strategy must be used. Given this known asymmetry in metacognitive performance between yes- and no-response (i.e., yes-response advantage) (Fleming and Dolan, 2010; Kanai et al., 2010) (Supplementary Figure 1), we here separately estimated the response-specific metacognitive sensitivity with a previously developed signal detection theoretic measurement (rs-meta-d’) (Maniscalco and Lau, 2014). rs-meta-d’ provides a bias-free measure of one’s ability to distinguish correct vs incorrect instances of a particular response type with confidence ratings. For instance, rs-meta-d’ for yes-responses measures an observer’s sensitivity for discriminating between correct yes-responses (hits) vs incorrect yes-responses (false alarms) by reporting higher confidence for the former than the latter.

We first conducted Experiment 1 where we behaviorally examined the metacognitive performance of schizophrenia patients and healthy controls in a visual detection task where the target could have either high or low SF (HSF or LSF). We then conducted Experiment 2 to examine the neural underpinnings of the group difference in metacognition with fMRI. Specifically, we examined whether the DLPFC, a region where metacognitive computations take place (Fleming et al., 2012a; Lau and Rosenthal, 2011), may show differential functional connectivity with the rest of the brain as a function of group, SF, and the response type (yes/no).

## 2. Experiment 1

In Experiment 1, schizophrenia patients and age matched controls performed a grating detection task and rated confidence in their detection responses. Gratings had either high or low SF (HSF and LSF, respectively). We measured metacognitive sensitivity separately for yes- and no-responses as a function of SF condition, and compared the effect of SF on metacognitive performance between the patient and control groups. We chose to use simple grating stimuli rather than more complex stimuli, such as faces, in order to avoid potential confounding variables due to other schizophrenia-related visual dysfunctions in processing complex stimuli (Revheim et al., 2014; Soria Bauser et al., 2012).

Perceptual sensitivity (d’) was prepared to be similar across groups and conditions by calibrating the grating contrast level prior to the detection task, which allowed us to selectively examine the potential differences in metacognitive performance.

### 2.1 Methods and Materials

#### 2.1.1 Participants

Twenty schizophrenia patients diagnosed with DSM-IV Axis I Disorders Patients Edition, Version 2.0 (SCID-P) (9 males, mean age = 42.3 ± 11.9) by psychiatrists at Kyoto University Graduate School of Medicine and eighteen healthy controls (12 males, mean age = 40.2 ± 11.6) were enrolled in this study. The data from three female patients were excluded from the analyses because of difficulty in comprehending the task, leaving data from the remaining seventeen patients for analysis (9 males, mean age = 40.5 ± 11.9). See supplementary Table 1 for their demographic information. The protocol was approved by the ethical committee of Kyoto University as well as by that of NICT. All participants gave written consent forms prior to participation.

#### 2.1.2 Procedure

Participants performed a pair of tasks to detect HSF (2.6 cycle per degree, cpd) and LSF (0.4 cpd) gratings (Figure 1A) in blocks, in a counterbalanced order. Each task consisted of a detection task block (360 trials, performed in six subblocks of 60 trials each), which followed two practice blocks (10 trials each) and one calibration block (80 trials). These two levels of SF were expected to evoke differential neural activity between schizophrenia patients and controls (Martinez et al., 2008). On every trial, a dynamic random noise pattern was displayed for 1.5 s. In half of these trials, after the first 1 s of random noise, a horizontal grating was superimposed on the noise pattern for 250 ms. Subsequently, participants were asked to indicate whether a grating had been presented or not (i.e., yes- or no-response) and then to rate their perceptual confidence on a four-point scale by using a keyboard.

**Figure 1.**
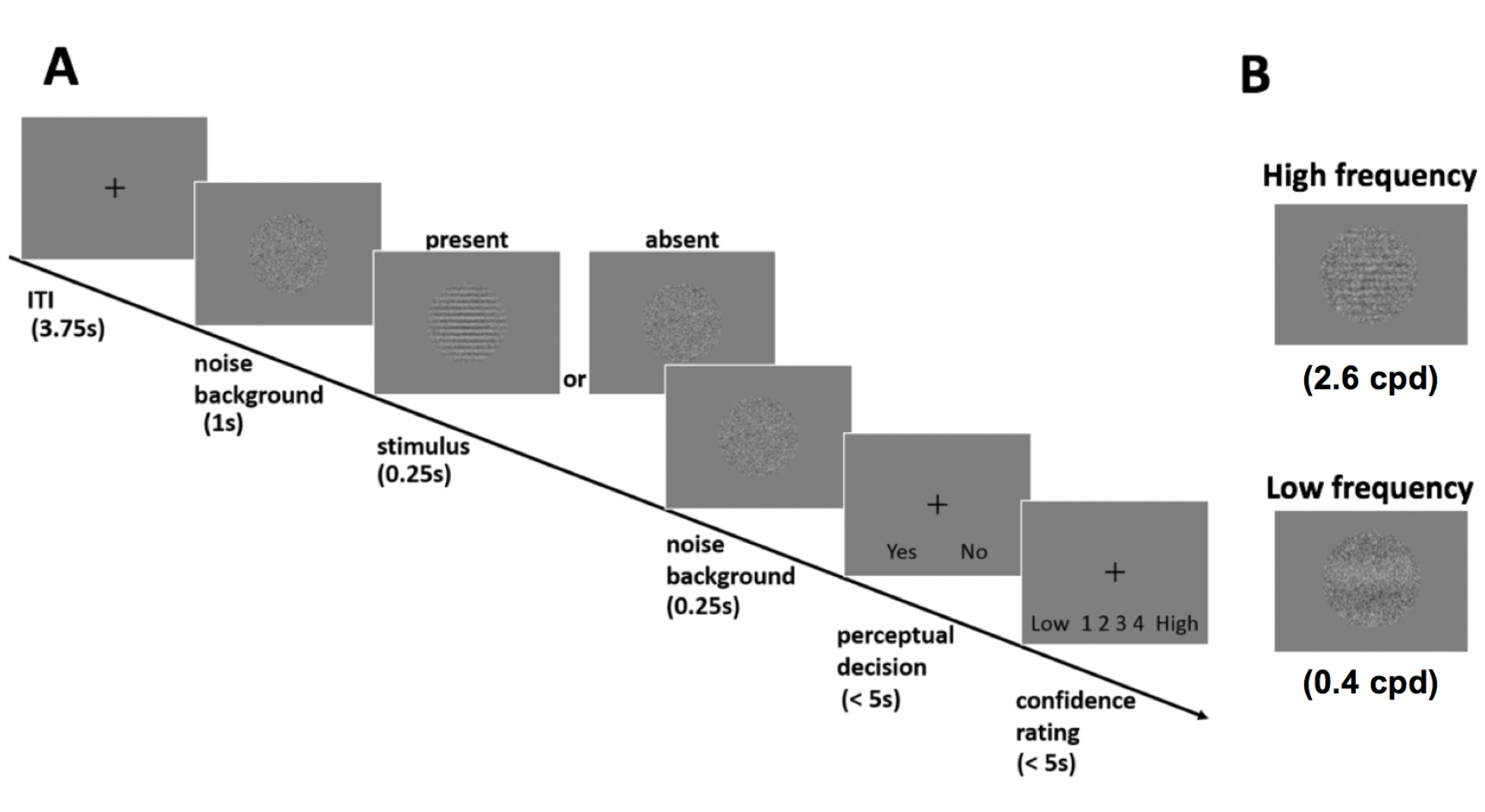
Schematics of the detection task. **A.** A trial sequence of the task. On half of the trials, a grating target briefly emerged and faded within a background patch containing dynamic white noise refreshed at 60 Hz (similar to TV static). Participants were asked to make a perceptual response indicating whether a grating was present (“yes” response) or absent (“no” response), and then to rate their confidence in their perceptual response with a four-responses scale. **B.** Two levels of spatial frequency of a grating target. A grating with high and low spatial frequency (2.6 cpd and 0.4 cpd, respectively) levels were used as a target in two separate sessions of Experiment 1 and 2. Grating phase was randomized across trials. ITI: inter-trial-interval. cpd: cycle per degree (in visual angle).

Prior to the task, the Michelson contrasts of the gratings were titrated with a QUEST threshold estimation procedure (Watson and Pelli, 1983) to achieve two slightly different levels of difficulty (hit rates of 55% and 65%). Calibration was conducted separately for each task with HSF or LSF gratings. Within each task, the level of difficulty was fixed within a given subblock and two levels of difficulty were alternated between subblocks. This alternation was intended to avoid over-habituation during the task by including slight variability of the stimuli. The data from all subblocks were used to estimate metacognitive performance for each task with HSF or LSF gratings in subsequent main analyses. The results are separately shown for each difficulty level in Supplementary Figure 4. See Supplementary material for further details.

#### 2.1.3 Analyses

Detection sensitivity was calculated as d’ for each SF level with standard signal detection theory (SDT) methods (Macmillan and Creelman, 1991). Metacognitive sensitivity for yes- and no-responses was separately calculated as response-specific meta-d’ (rs-meta-d’; http://www.columbia.edu/~bsm2105/type2sdt/fit_rs_meta_d_MLE.m) (Maniscalco and Lau, 2014) for each SF level. High values of rs-meta-d’ for yes-responses indicate that the participant’s confidence ratings discriminate well between correct and incorrect yes-responses, i.e. mean confidence is higher for hits than for false alarms. Similarly, high values of rs-meta-d’ for no-responses indicate that the participant’s confidence ratings discriminate well between correct and incorrect no-responses, i.e. mean confidence is higher for correct rejections than for misses. Here, the ratio of meta-d’ (metacognitive sensitivity) to d’ (perceptual sensitivity) (meta-d’/d’) was then calculated for each SF level to quantify metacognitive efficiency (Fleming and Lau, 2014; Maniscalco and Lau, 2014). Meta-d’ corresponds to the degree of metacognitive sensitivity (to discriminate correct versus incorrect perceptual response), which is expected to equal to the value of perceptual sensitivity d’ (to discriminate stimulus presence or absence with yes- or no-response, respectively) when the signal strength in SDT framework is equally contributing to d’ and meta-d’. Thus, for a subject whose behavior matches SDT expectation, meta-d’ = d’, meaning that the subject’s measured metacognitive sensitivity (meta-d’) corresponds to the level of metacognitive sensitivity that would be expected to arise from their performance on the primary perceptual task (d’). For subjects with meta-d’ < d’, metacognitive sensitivity is lower than would be expected based on primary task performance. For such subjects, metacognitive sensitivity can be considered to be “inefficient” or “suboptimal,” relative to SDT expectation.

Across-participant means for d’, meta-d’, and meta-d’/d’ were analyzed with Analysis of Variance (ANOVA) in SPSS version 25 (IBM). For subsequent analyses, an ANOVA with two within-participant factors of Response-type (yes/no) and SF (high/low) as well as with one between-participant factor of a Group were conducted.

### 2.2 Results and discussion

First, the analysis of Meta-d’/d’ revealed a main effect of Response-type (F(1, 33) = 25.99, p < .001, partial η2 = .441) (Figure 2). This main effect was due to higher Meta-d’/d’ following yes-than no-responses, which is consistent with previous studies (Fleming and Dolan, 2010; Kanai et al., 2010) and our pilot experiment (Supplementary Material). Interestingly, such yes-response advantage was modulated by Group and SF (a three-way interaction: F(1, 33) = 4.60, p = .039, partial η2 = .122). Post-hoc t-tests revealed that controls showed significantly higher Meta-d’/d’ following yes-than no-responses (i.e., yes-response advantage) only when detecting an HSF grating target (p < .001) but not LSF grating target (p = .169). By comparison, patients showed significant yes-response advantage irrespective of target SF (HSF: p = .013; LSF: p = .001). There was no significant main effect of Group (F(1, 33) = .77, p = .386), suggesting that patients did not show any generic reduction of metacognitive performance relative to controls. The magnitude of yes-response advantage (i.e., the difference in meta-d’/d’ between yes- and no-responses, Figure 2B) was significantly larger with HSF than LSF targets among controls (t(17) = 2.22, p = .040) but not among patients (t(16) = −.63, p = .540). The degree to which confidence rating captured the accuracy of perceptual response qualitatively mirrored the results of Meta-d’/d’ (Supplementary Figure 2).

**Figure 2.**
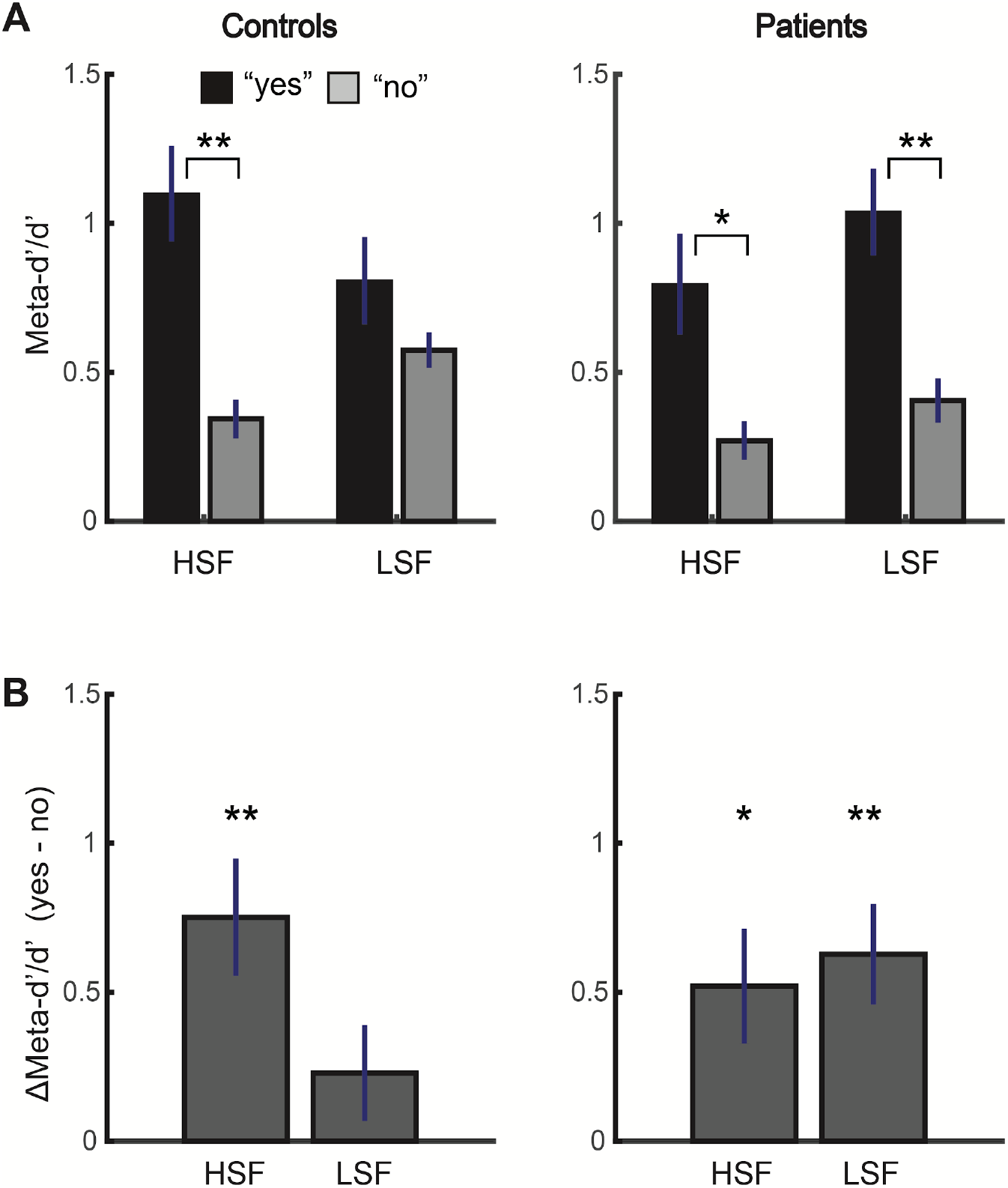
Differences in Meta-d’ between schizophrenia patients and controls as a function of response type (yes/no) and that of spatial-frequency (HSF/LSF) in Experiment 1. **A.** Controls showed advantageous metacognitive performance with yes-relative to no-responses only during the HSF target detection task but not during the LSF target detection task, whereas patients showed advantageous performance with yes-responses irrespective of the target spatial-frequency. There was a significant spatial-frequency x response type x group interaction (F(1, 33) = 4.6, p = .039). The results of post-hoc t-tests are shown. **B.** Same result from the analyses depicted in A. Differences in meta-d’/d’ between yes- and no-responses are shown for demonstrative purposes. Here, larger values indicate more advantageous metacognitive performance with yes-than no-responses. Error bars indicate standard error of the mean. ** p<.01, * < p<.05.

For the detection sensitivity d’ (Supplementary Figure 3), there was neither a significant main effect of Group (F(1, 33) = .16, p = .694) nor a significant interaction between Group and SF (F(1,33) = 3.80, p = .060), suggesting that patients did not statistically differ from controls in their perceptual performance. There was a main effect of SF (F(1, 33) = 4.33, p = .045, partial η2 = .116), which was due to lower d’ with LSF relative to HSF stimuli (see Supplementary Material). A potential relationship between hallucination severity (measured with Positive and Negative Syndrome Scale (PANSS) (Kay et al., 1987; Kay et al., 1991)) and metacognitive performance among patients, although tentative, can be found in Supplementary Material.

Taken together, the results suggest that insensitivity to SF modulation may characterize metacognitive performance among patients (Figure 2).

## 3. Experiment 2

Experiment 1 indicated that only controls show SF dependency in their metacognitive performance, while patients do not. We predicted that controls, but not patients, may rely less on the neural circuit typically supporting metacognition with LSF stimuli. A large body of literature has shown that DLPFC typically plays a central role in metacognition (Cortese et al., 2016; Fleming et al., 2012a; Fleming et al., 2012b; Koizumi et al., 2016; Lau and Passingham, 2006; Rounis et al., 2010). However, whether DLPFC atypically contributes to metacognition among schizophrenia patients remains unknown. We here particularly predicted that DLPFC may be a potential source of the group difference in metacognitive function. To test this, we next conducted the same experimental task as Experiment 1 in an fMRI scanner.

### 3.1 Methods and Materials

#### 3.1.1 Participants

Seventeen schizophrenia patients diagnosed with DSM-IV Axis I Disorders Patients Edition, Version 2.0 (SCID-P) (6 males, mean age = 42.8 ± 9.5) and seventeen healthy controls (7 males, mean age = 40.2 ± 10.1) were enrolled in this study. The data from two male patients were excluded from the analyses because of their inability to respond within the limited windows (< 2 s). The data from the remaining fifteen patients (4 males, mean age = 44.7 ± 8.4) were analyzed (see Supplementary Table 1). The protocol was approved by the ethical committee of Kyoto University as well as by that of NICT. All participants gave written consent forms prior to participation.

#### 3.1.2 Procedure

The task procedure was similar to that of Experiment 1. The runs were blocked for HSF and LSF stimuli, and the order of SF level was counterbalanced across participants. Participants viewed the stimuli presented on a monitor screen via a mirror, and performed the task with an MRI compatible response keypad (Current Designs, Inc.). See Supplementary Material for more details.

#### 3.1.3 ROI definition

The DLPFC ROI was functionally defined. Specifically, the clusters of voxels that were significantly activated for the confidence rating relative to the fixation period from all the trials (p < .01, Bonferroni-corrected) were selected, which were located within Brodmann area (BA) 46 (MRIcron, Brodmann 48 Area Atlas Template, https://people.cas.sc.edu/rorden/mricron/index.html) (Figure 3). For the DLPFC ROI, the center of gravity in Talairach space was [X = −34.28 (± SD 2.45), Y = 35.33 (± 3.37), Z = 32.43 (± 3.81)] for the left hemisphere cluster and [X = 37.67 (± 3.52), Y = 30.44 (± 3.43), Z = 31.61 (± 3.35)] for the right hemisphere cluster.

**Figure 3.**
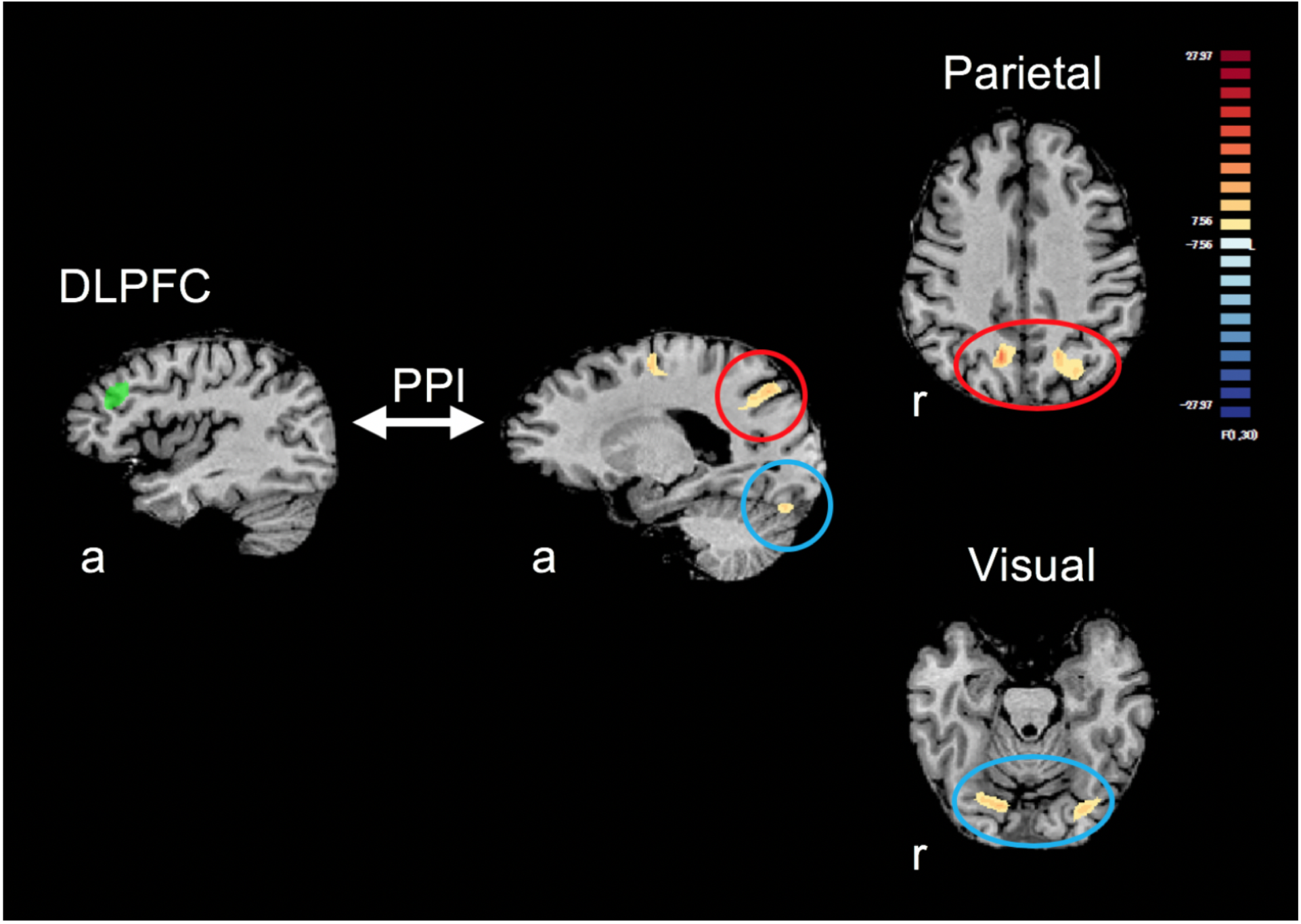
The results of a whole-brain analysis examining where in the brain showed altered functional connectivity with the DLPFC during confidence rating as a function of spatial-frequency, response-type, and group. Here, the degree of functional connectivity was estimated with a general form of context-dependent psychophysiological interaction (gPPI) (McLaren et al., 2012). The functional connectivity between the DLPFC and bilateral clusters in parietal and visual cortices were significantly modulated as a function of the interaction between spatial-frequency, response-type, and group (p < 0.01, corrected with cluster-size thresholding). Here, DLPFC ROIs were functionally defined from a group GLM. ROIs include the voxels that showed significantly larger activity during the confidence rating period relative to fixation in a group GLM (p < .01, Bonferroni corrected). a: anterior, r: right, PPI: psychophysiological interaction.

As a control ROI which is related to visual processing but not directly to metacognitive processing (Cortese et al., 2016), the clusters of voxels that were significantly activated for the stimulus period relative to the fixation from all the trials (p < .01, Bonferroni-corrected) were selected within visual cortex (Supplementary Figure 7A). For the control ROI within visual cortex, the center of gravity in Talairach space was [X = −22.21 (± SD 3.88), Y = −81.09 (± 5.47), Z = −10.29 (± 5.03)] for the left hemisphere and [X = 22.40 (± 5.96), Y = −81.35 (± 5.56), Z = −8.27 (± 2.78)] for the right hemisphere.

#### 3.1.4 Analyses

The fMRI data were preprocessed and analyzed with BrainVoyager 21.0 (Brain Innovation, the Netherlands). We aimed to examine whether functional connectivity between DLPFC and other brain areas was modulated as a function of SF, Response-type, and Group. With this aim, we first conducted a whole-brain Generalized form of context-dependent psychophysiological interaction (gPPI) analysis (McLaren et al., 2012) with DLPFC as a seed ROI. This allowed us to quantify the functional integration of DLPFC with other brain areas as a function of the interaction between group, SF and response-type at the participant level.

In addition, to decode the trial-by-trial confidence level in a binary manner, high or low confidence, from the multivoxel activation pattern in DLPFC during the confidence rating period (i.e., predicting the confidence level on each trial from the activation pattern), we built a decoder with sparse logistic regression (SLR) which automatically selects relevant features (i.e., voxels) (Yamashita et al., 2008).

### 3.2 Results and discussion

#### 3.2.1 Behavioral results

The behavioral data were not of main interest in Experiment 2, given that there were too small trial numbers in Experiment 2 (96 trials) relative to Experiment 1 (360 trials) to properly fit the data to estimate Meta-d’. The overall results were qualitatively similar to that of Experiment 1 (Supplementary Figure 5).

#### 3.2.2 fMRI results

The whole-brain analysis of gPPI effect revealed that the DLPFC ROI showed altered functional connectivity as a function of SF, Response-type, and Group with clusters in parietal and visual cortices (p < .01, corrected with cluster-level thresholding (Forman et al., 1995; Goebel et al., 2006)) (Figure 3). Note that here, the parietal and visual cortices were not predetermined as ROIs but were shown significant in the whole brain analysis. The bilateral clusters in the parietal cortex and visual cortex were located posterior of BA7 and ventral of BA18 (MRIcron, Brodmann 48 Area Atlas Template), respectively (see Supplementary Material for the center of gravity in Talairach space for each ROI). The centers of gravity for the left and right clusters in parietal cortex were [X = −21.34 (± 4.76), Y = −68.37 (± 4.18), Z = 38.32 (± 3.40)] and [X = 10.50 (± 2.28), Y = −61.86 (± 3.13), Z = 36.32 (± 2.41)], respectively. The centers of gravity for the left and right clusters in visual cortex were [X = 17.13 (± 4.62), Y = −72.81 (± 2.71), Z = −16.73 (± 2.92)] and [X = −24.0 (± 5.21), Y = −77.51 (± 3.22), Z = −19.52 (± 1.93)], respectively.

For illustrative purposes only, the parameter estimates (Beta) for PPI terms in the parietal and occipital clusters (averaged between the hemispheres) are visualized separately for each level of SF, response-type, and group in Supplementary Figure 6. This result, although illustrative, shows that the functional connectivity between DLPFC and parietal or visual cortex was more enhanced following no-relative to yes-responses with the HSF stimuli among controls. Considering that controls behaviorally show yes-response advantage in metacognitive performance only with the HSF stimuli (Experiment 1), one interpretation of this functional integration effect may be that there was more effortful (thus enhanced) integration of sensory and decision information by the DLPFC following no-response to the HSF stimuli.

The results of the multivoxel decoding analysis also supported the possibility that patients and controls may differ in DLPFC recruitment during metacognitive judgement (Figure 4). That is, the decoding analysis with the DLPFC ROI revealed a significant interaction between Group and SF (F(1, 27) = 5.08, p = .032, partial η2 = .158). This interaction was due to higher decoding accuracy for HSF relative to LSF stimuli among controls (p = .043, partial η2 = .144), while there was no such SF dependence of decoding accuracy among patients (p = .289). There was no significant main effect of Group (F(1, 27) = .07, p = .799) or SF (F(1, 27) = 2.81, p = .105). Unlike with DLPFC, there was no significant difference in decoding accuracy as a function of Group and SF within the control ROI in a visual cortex (Supplementary Figure 7B). There was no significant main effect of Group (F(1, 27) = 2.33, p = .138) or SF (F(1, 27) = 0.10, p = .751), and their interaction was also not significant (F(1, 27) = 2.15, p = .154). Note that the factor of Response-type was not considered in the decoding analysis due to the limits in trial number (see *Decoding analysis* in Supplementary Material).

**Figure 4.**
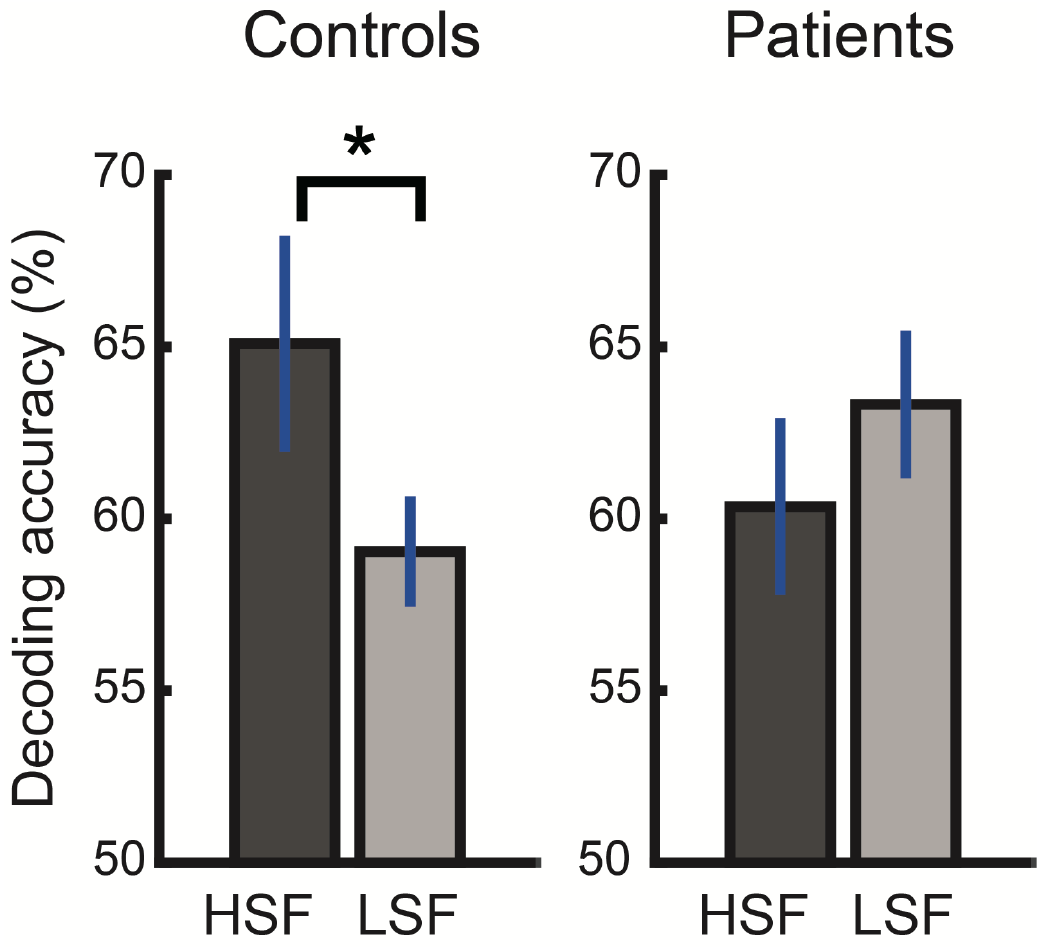
There was a significant interaction between group and spatial frequency in the decoding accuracy of the trial-wise confidence level (high/low) from the multivariate activation patterns in the DLPFC ROI (shown in Figure 3). Only among controls, decoding accuracy was significantly higher with HSF than LSF stimulus judgement, whereas it was statistically similar between the spatial-frequency levels among patients. a: anterior, r: right, * p < .05.

Consistent with Experiment 1, patients and controls did not differ in their overall behavioral performance in metacognition and perception (i.e., unless considering response-type), regardless of the SF level (Supplementary Material). Thus, it is unlikely that the decoding accuracy difference merely reflected the metacognitive performance difference. Instead, such results more likely reflect the group difference in the involvement of DLPFC in metacognition, as a function of SF.

## 4. General discussion

While a potential relationship between schizophrenia and atypical metacognitive function has been speculated for decades (Frith and Done, 1988), whether schizophrenia patients have altered metacognitive ability to introspect perception has remained inconclusive (Charles et al., 2017; Powers et al., 2017; Rouault et al., 2018). We here show that patients and controls generally perform equally well in a visual metacognitive task. However, the groups differed in both behavioral performance and neural activity in terms of their dependency on SF.

Overall, metacognitive efficiency for introspecting perceptual accuracy was higher for yes-responses (i.e., perception of target presence) than for no-responses (i.e., absence), consistent with earlier studies (Fleming and Dolan, 2010; Kanai et al., 2010). A novel finding of this study was that patients and controls differ in the SF range where they display the yes-response advantage in metacognition. Specifically, controls showed a yes-response advantage only when detecting a higher SF visual target but not when detecting a lower SF target. Meanwhile, patients showed a yes-response advantage irrespective of the SF level.

An fMRI study (Experiment 2) further supported such SF independence of metacognitive function among patients. That is, the functional connectivity between DLPFC and parietal or visual cortex during the metacognitive judgement was differently modulated as a function of the group, SF, and response type. Moreover, among controls, trial-by-trial confidence level could be more accurately decoded from the multivoxel patterns in DLPFC with higher than lower SF stimuli. Meanwhile, decoding accuracy was similar between the SF levels among patients.

To provide explanations for the group difference in SF dependence, we first turn to why there is generally a yes-response advantage in metacognition. It has been repeatedly demonstrated that metacognitive judgements are mainly based on perceptual evidence that contributed to finalize perceptual decisions (Koizumi et al., 2015; Maniscalco et al., 2016; Peters et al, 2017; Zylberberg et al., 2012). While these findings are mainly based on discrimination tasks, a similar heuristic mechanism may be at play with a detection task. In a detection task, there is perceptual evidence to support yes-responses (stimulus presence) but no evidence to support no-responses (stimulus absence). Thus, perceptual confidence for only the yes-responses likely reflects the amount of supporting evidence, which is typically larger for correct than incorrect responses. However, confidence for no-responses could be more difficult to estimate, because supporting evidence for no-responses is absence of evidence, which might provide a poorer basis for discerning correct versus incorrect responses. The yes-response advantage in a given perceptual condition may be considered as an index for the degree to which supporting perceptual evidence is directly translated to metacognitive judgements (i.e., confidence rating).

Considering this, one possibility is that the group difference in yes-response advantage reflects the degree to which metacognitive judgement is based on perceptual evidence in a given SF range. Individuals *without* schizophrenia may base their metacognitive judgements more on higher SF evidence because it typically contributes more to finalized perceptual decisions (Bar et al., 2006). Meanwhile, schizophrenia patients may also rely on evidence in the lower SF range, potentially because it tends to contribute more to finalized perceptual decisions relative to controls (Laprevote et al., 2010; Laprevote et al., 2013).

Previous studies have shown reduced neural responses and lowered perceptual sensitivity to low SF stimuli among schizophrenia patients (Butler and Javitt, 2005; Martinez et al., 2012; Martinez et al., 2008; Slaghuis and Curran, 1999) although see also (Keri et al., 2002). The current results, together with the findings at the early visual stages, suggest the possibility that the early stage sensory inputs across SF ranges may get wired to higher-order metacognitive function differently among schizophrenia patients compared with controls.

The fMRI results (Experiment 2) suggested that the group difference in metacognition may be at least partly related to the function in the DLPFC. First, the result of gPPI analysis showed that controls and patients differ in how the functional connectivity between the DLPFC and parietal or visual cortex is modulated during metacognitive judgement, as a function of the stimulus SF and the response type (yes/no) (Figure 3). Given that parietal cortex and visual cortex are each involved in perceptual decisions (Kiani and Shadlen, 2009) and sensory evidence processing (Cortese et al., 2016), the result suggests that the degree to which the DLPFC engages to integrate the perceptual-level information in parietal and visual cortices differs between the groups, depending on whether they perceived stimulus presence (yes- or no-responses) and on whether the stimulus was high or low in its SF. Whether the degree of such functional connectivity reflects more effortful metacognitive processing (Supplementary Figure 6) or other computational differences remain to be further examined in future studies with larger sample size.

The decoding analysis results further supported the possibility that DLPFC may at least partly contribute to group difference in metacognition. The decoding results showed that, among controls, metacognitive judgements (i.e., trial-wise confidence ratings) are more accurately decoded from the activation patterns in DLPFC during the detection of high SF stimuli than that of low SF stimuli. Meanwhile, patients did not show such SF dependency. While controls may rely on the DLPFC when estimating perceptual confidence in a SF-specific manner, schizophrenia patients may rely on the DLPFC in a more uniform manner across the SF range.

What remains to be explained is the result that metacognitive efficiency for no-responses is generally lower than yes-responses but is not completely abolished (i.e., above-zero Meta-d’). This suggests that participants (both controls and patients) may rely on some compensatory strategies to evaluate their perceptual confidence for no-responses, which are alternative to, but not mutually exclusive with, the aforementioned strategy to rely on sensory evidence that contributed to perceptual decisions (Koizumi et al., 2015; Maniscalco et al., 2016; Peters et al, 2017; Zylberberg et al., 2012). One such alternative strategy could be to rely on the level of attention allocated on given trials (Kanai et al., 2010). As attentional level is expected to be generally higher for correct than for incorrect responses, such a difference in attentional level can be mapped onto higher confidence for correct responses, resulting in some degree of metacognitive sensitivity. This strategy based on attentional level difference may also partly contribute to metacognitive performance with LSF stimuli among controls, where metacognitive efficiency was similar between yes- and no-responses. If participants were to evaluate their perceptual confidence for correct versus incorrect responses based on the attentional level difference between those responses, then metacognitive sensitivity is expected to be similar between yes- and no-responses. This is because attentional level allocated to the task stimuli is expected to be similar between stimulus present and absent trials (which, other things being equal, would map onto correct yes- and no-responses when attentional level is sufficiently higher), given that participants had no prior information as regards to which trial (i.e., target present or absent) will be presented. Schizophrenia patients, on the other hand, may have less flexibility to switch between different metacognitive strategies across SF ranges.

The result showed that metacognitive efficiency for no-responses differs between the tasks with HSF and LSF stimuli among controls, despite the fact that stimulus absent trials (which constitute the ‘correct’ no-responses) were identical between the tasks. This result likely owes to a general limitation in perceptual decision making strategy which results in a failure to alternate between different perceptual strategies across trials when the trials are performed in a single block (Rahnev et al., 2010; Gorea and Sagi, 2000). That is, controls are likely to evaluate perceptual confidence with one strategy uniformly within a given task, depending on SF of the target stimuli. And the use of different strategies across the tasks may explain the difference in metacognitive efficiency for no-responses between the tasks with different SF levels. Future studies may further examine this possibility with a non-blocked design where both HSF and LSF stimuli are presented as targets within a single block.

Lastly, whether patients show atypicality in metacognitive performance with other sensory modalities such as audition remains one important question to be examined, as hallucinations among schizophrenia patients are even more prevalent in the auditory domain than the visual domain (59% and 27% prevalence, respectively) (Waters et al., 2014). As a previous study demonstrated that one’s metacognitive performance correlates across the sensory modalities (Faivre et al., 2018), future studies may examine whether metacognitive function among patients also show atypical dependence on some features of auditory stimuli.

## 5. Conclusions

Taken together, our results demonstrate that schizophrenia patients do not show SF dependence in their metacognitive performance, while controls show differential metacognitive performance as well as differential reliance on DLPFC across the SF levels. The finding among controls is itself novel and may suggest that they can flexibly shift their metacognitive strategy depending on stimulus features. Inability of patients to adaptively shift their metacognitive strategy across visual conditions might contribute to their altered subjective experience of the sensory world.

## 6. Acknowledgements

We thank Aurelio Cortese for his helpful comments on the analysis and Raihyung Lee, Ben Smith, Matthias Michel, Kiyofumi Miyoshi for their helpful comments on the manuscript. This work was supported by JSPS KAKENHI Grant-in-Aid for Scientific Research B [19H03583 to HT, 18H02714 to AK, 17H04248 to JM], MEXT Grant-in-Aid for Scientific Research on Innovative Areas [16H06572 to HT, 18H05130 to JM], and the Strategic Research Program for Brain Sciences by the Japan Agency for Medical Research and Development (AMED) [19dm0307008h0002 and 19dm0107151s0104 to HT], JST presto awarded each to AK and KA. The funders played no scientific role in the project.

