## Supplementary material for "Atypical spatial frequency dependence of visual metacognition among schizophrenia patients"

### Supplementary Materials

**Supplementary Table 1:** Demographic information on patients and controls enrolled in Experiment 1 and 2.

| | | Age | Gender | FSIQ | chlorpromazine equivalent | PANSS positive score | PANSS negative score | PANSS general psychopathological | Years of education | Employment or education |
|---|---|---|---|---|---|---|---|---|---|---|
| | | Mean (SD) | Male / Female | Mean (SD) | Mean in mg (SD) | Mean (SD) | Mean (SD) | Mean (SD) | Mean (SD) | Full / Part / None |
| **Experiment 1** | **Patients** | 40.5 (11.9) | 9 / 8 | 108.2 (7.8) | 547.4 (455.6) | 15.9 (5.5) | 20.6 (2.6) | 35.9 (7.5) | 14.1 (2.3) | 2 / 8 / 7 |
| | **Controls** | 40.2 (11.6) | 12 / 6 | 112 (5.0) | N/A | N/A | N/A | N/A | 16.9 (1.5) | 18 / 0 / 0 |
| **Experiment 2** | **Patients** | 44.7 (8.4) | 4 / 11 | 98.9 (9.0) | 686.3 (563.1) | 15.6 (5.2) | 19.0 (4.9) | 33.4 (10.3) | 14.3 (1.6) | 2 / 4 / 9 |
| | **Controls** | 40.2 (10.1) | 7 / 10 | 110.5 (5.7) | N/A | N/A | N/A | N/A | 16.6 (1.1) | 16 / 1 / 0 |

*FSIQ: Full scale intelligence quotient, PANSS: Positive and Negative Syndrome Scale

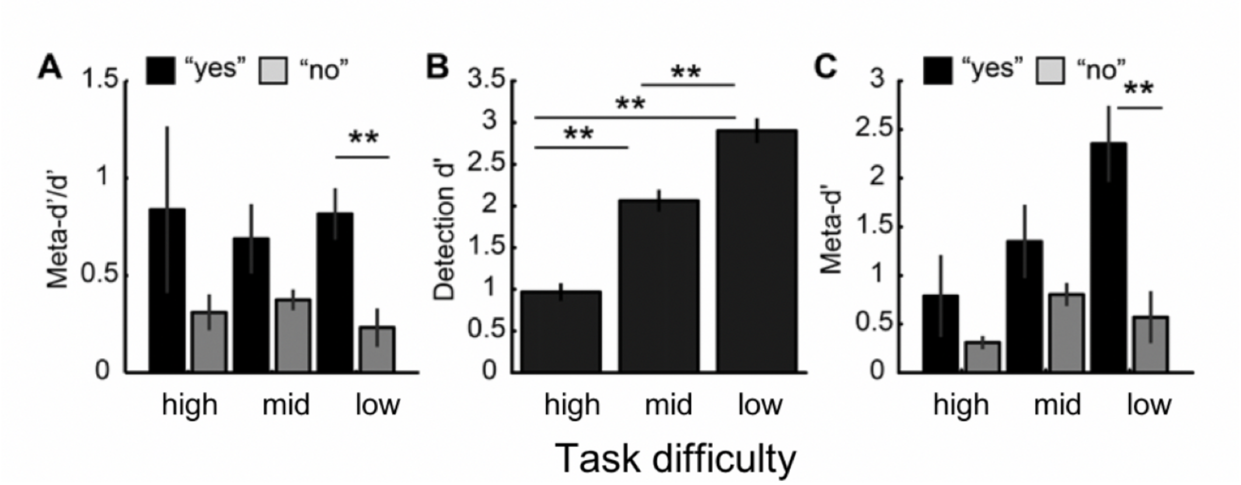


**Supplementary Figure 1.** Metacognitive and perceptual sensitivity as a function of task-difficulty level among healthy participants in Pilot Experiment. **A.** Metacognitive performance is quantified as Meta-d'/d' to depict metacognitive sensitivity (Meta-d') in relation to perceptual sensitivity (d'). Meta-d'/d' was numerically higher following "yes" than "no" responses in general, although a main effect of Response-type was not significant (F(1, 14) = 3.707, p =. 075). There was a significant interaction between Response-type and Task-difficulty (F(2, 13) = 4.268, p =.038), which was due to that there was a significant difference between response types only with low task difficulty level (p = .008) but not with mid and high difficulty levels (p = .138, p = .217, respectively). **B.** The across-participant means for Detection *d'* showing perceptual sensitivity during the detection task, which monotonically decreased with task difficulty. **C.** The across-participant means for response-specific Meta-d'. Similarly to Meta-d'/d' shown in A, there was a significant interaction between Response-type and Task-difficulty (F(2, 13) = 6.716, p =.01). This interaction was due to that yes-response advantage was significant only with low task-difficulty level (i.e., highest contrast) (p = .006) but not with mid and high difficulty levels (p = .225, p = .276, respectively). Error bars indicate standard error of the mean. ** *p*<.01.

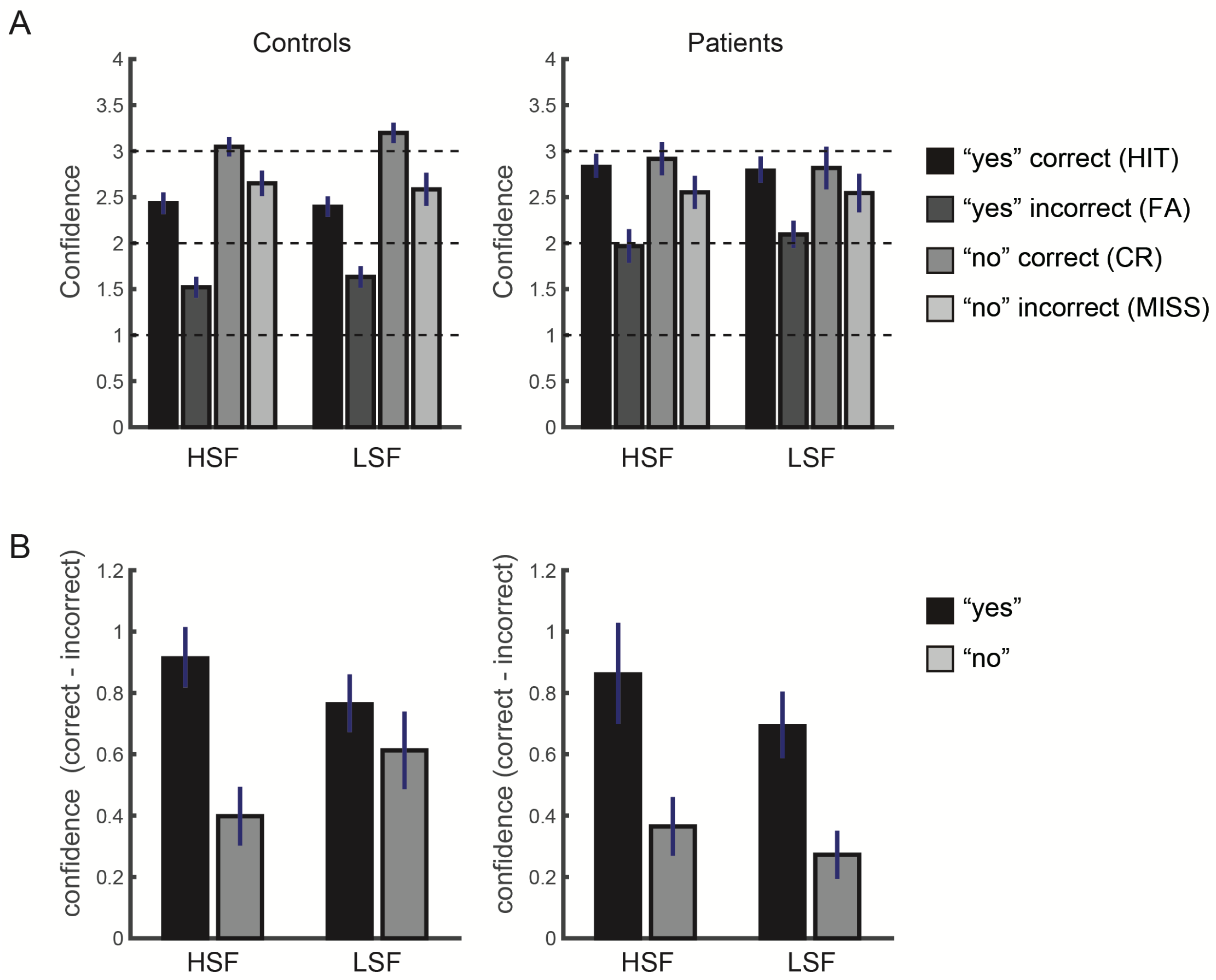


**Supplementary Figure 2.** Demonstration of confidence rating on correct and incorrect trials in Experiment 1. **A.** Mean confidence rating for each trial type, correct "yes" responses (i.e., Hit), incorrect "yes" responses (i.e., False Alarm, FA), correct "no" responses (i.e., Correct Rejection, CR), and incorrect "no" responses (i.e., Miss). Means are plotted separately for HSF and LSF conditions. **B.** Mean difference in confidence rating between correct and incorrect responses per response type (i.e., yes- and no-responses). Means are plotted separately for HSF and LSF conditions. Error bars indicate standard error of the mean. HSF, High spatial-frequency, LSF, Low spatial-frequency. *Related to Figure 2.*

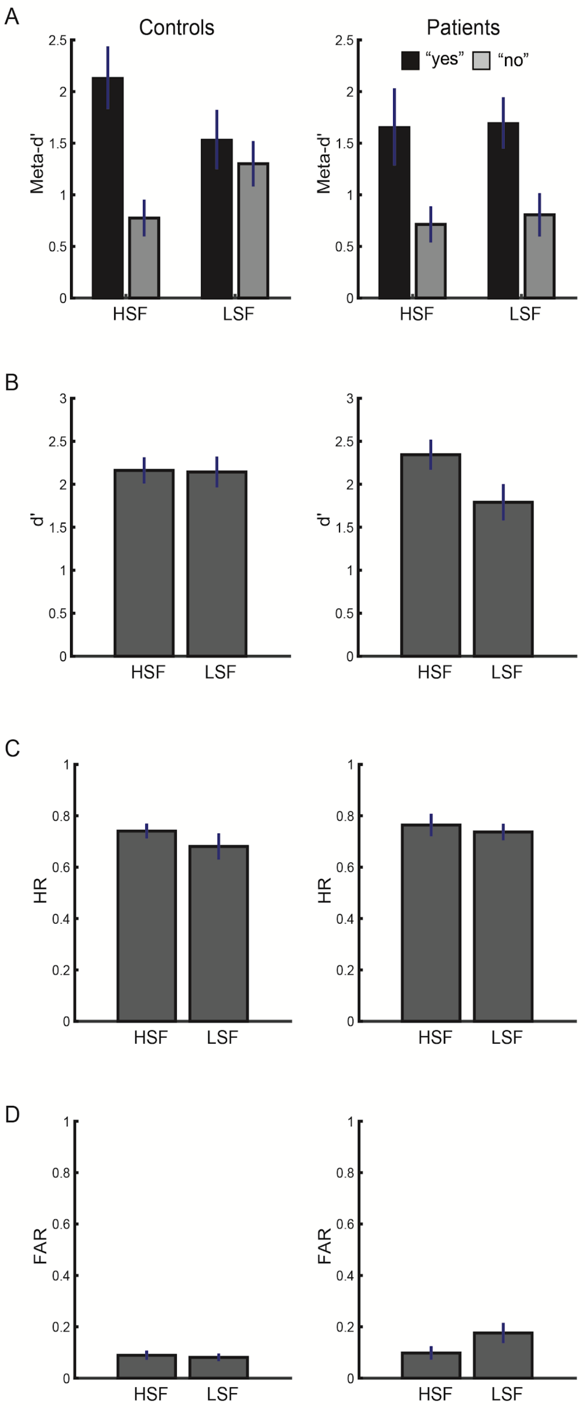


**Supplementary Figure 3.** The results of meta-d', d', HR and FAR in Experiment 1. **A.** The result of meta-d' qualitatively mirrored that of meta-d'/d', although a three-way interaction did not reach significance ($F(1, 33) = 3.38$, $p = .075$). There was a significant main effect of Response-type ($F(1, 33) = 16.11$, $p < .000$), also replicating the general yes-response advantage in metacognition. **B.** For the detection sensitivity d', there was neither a significant main effect of Group ($F(1, 33) = .16$, $p = .694$) nor a significant interaction between Group and Spatial-frequency ($F(1,33) = 3.80$, $p = .060$), suggesting that Patients did not statistically differ from Controls in their perceptual performance. There was a main effect of Spatial-frequency ($F(1, 33) = 4.33$, $p = .045$, partial $\eta 2 = .116$), which was due to generally lower d' with LSF relative to HSF stimuli. **C.** For hit rate (HR), there was no significant difference in HR as a function of Spatial-frequency ($F(1, 33) = 1.80$, $p = .188$), Group ($F(1, 33) = .73$, $p = .40$), and their interaction ($F(1, 33) = .26$, p =. 612). **D.** Meanwhile, for the analysis of false alarm rate (FAR), there was a significant interaction between Spatial-frequency and Group ($F(1, 33) = 5.67$, $p = .023$, partial $\eta 2 = .147$). The main effect of Spatial-frequency and that of Group were not significant ($F(1, 33) = 3.71$, $p = .063$; $F(1, 33) = .59$, $p = .117$, respectively). Post-hoc analyses showed that this was due to the fact that there was significant difference in FAR between HSF and LSF only among Patients ($p = .005$) but not among Controls ($p = .746$). The FAR was significantly higher for Patients than Controls, which was the case with LSF ($p = .029$) but not with HSF ($p = .789$). This result of FAR suggests that visual processing is atypically modulated by spatial-frequency among Patients. Note that this difference in FAR cannot explain away the results of metacognitive performance difference described in the main text. If we were to expect that differences in FAR account for that in metacognition (e.g., Meta-d'/d'), we would have observed the difference in metacognitive performance between HSF and LSF among Patients. However, this possibility is unlikely as it is contrary to the actual results (Figure 2). Error bars indicate standard error of the mean. HSF, High spatial-frequency, LSF, Low spatial-frequency. *Related to Figure 2.*

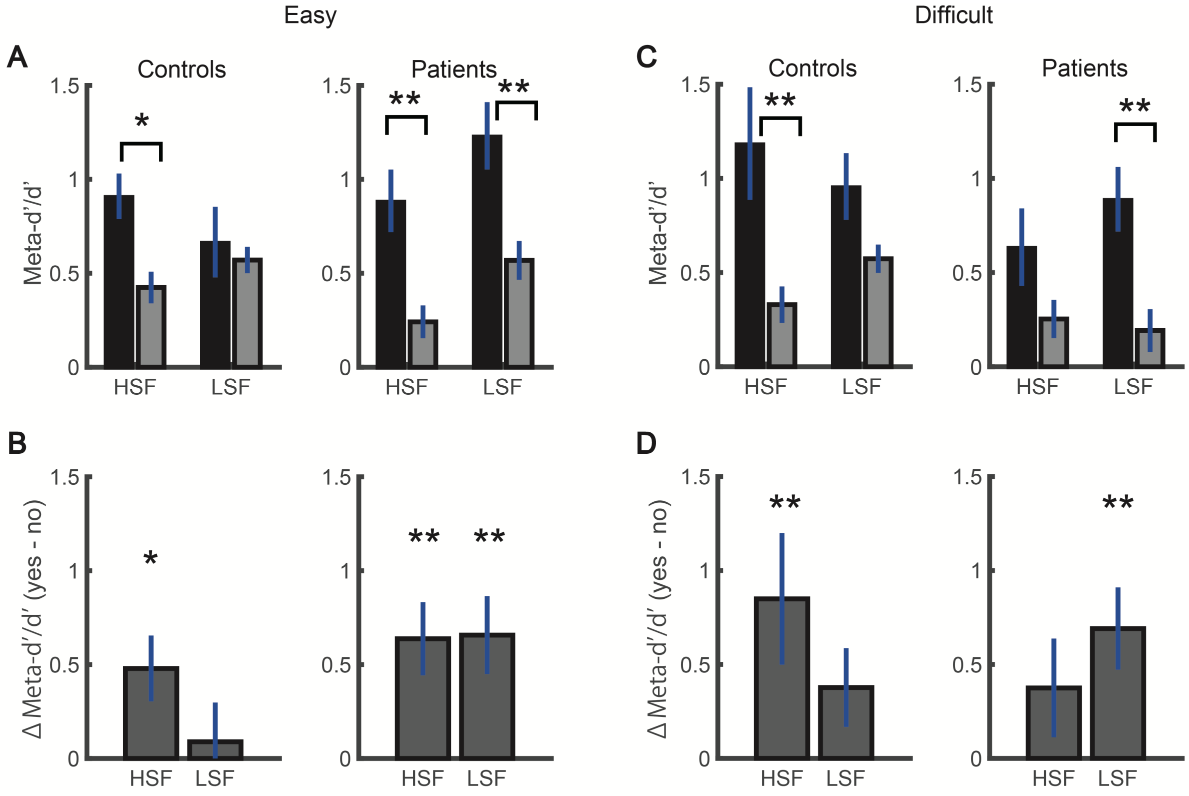


**Supplementary Figure 4.** Differences in Meta-d' between schizophrenia patients and controls as a function of response type (yes/no) and that of spatial-frequency (HSF/LSF) in Experiment 1. Here, metacognitive efficiency (Meta-d'/d') in Experiment 1 is separately estimated for the two levels of difficulty, whereas the data from both difficulty levels were merged to estimate metacognitive efficiency in the main text to ensure more stable estimations of efficiency (Figure 2). **A.** In an easy condition (i.e., higher stimulus contrast), controls showed advantageous metacognitive performance with yes- relative to no-responses only during the HSF target detection task but not during the LSF target detection task, whereas patients showed advantageous performance with yes-responses irrespective of the target spatial-frequency. Not surprisingly given a relatively small trial numbers used for the estimation of Meta-d'/d' (180 trials, which is a half of the trials included in the estimates shown in Figure 2), there was no significant spatial-frequency x response type x group interaction ($F(1, 33) = 1.27$, $p = .268$). The results of post-hoc t-tests are shown. **B.** Same result from the analyses depicted in A. Differences in meta-d’/d' between yes- and no-responses are shown for demonstrative purposes. Here, larger values indicate more advantageous metacognitive performance with yes- than no-responses. **C.** In a difficult condition (i.e., lower stimulus contrast), controls showed yes-response advantage only during the HSF target detection task but not during the LSF target detection task, whereas patients showed yes-response advantage only during the LSF target detection task but not during the HSF target detection task. There was no significant spatial-frequency x response type x group interaction ($F(1,33) = 2.41$, $p = .130$). **D.** Same result from the analyses depicted in C. Error bars indicate standard error of the mean. ** $p<.01$, * < $p<.05$.

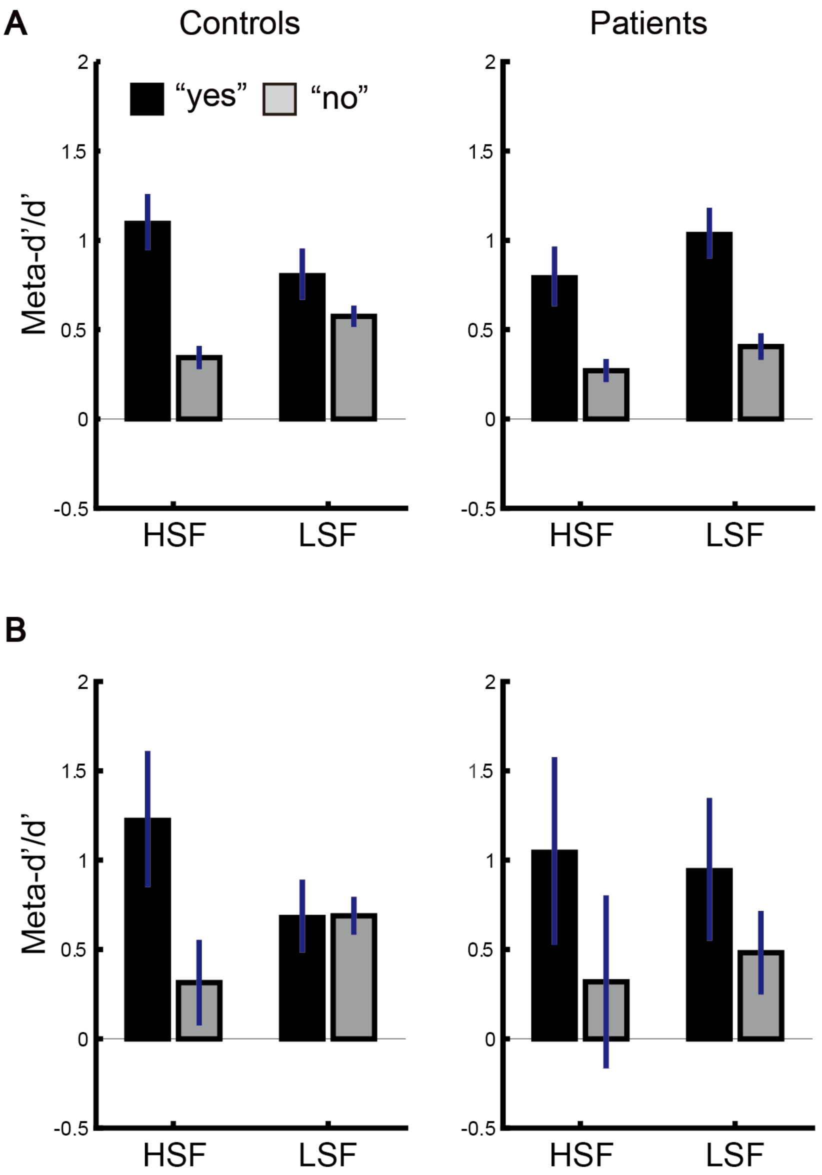


**Supplementary Figure 5.** Comparison of behavioral results from Experiment 1 (**A**) and 2 (**B**). **A.** The results reproduced from Experiment 1 presented in Figure 2. **B.** The results of meta-d'/d' from Experiment 2 qualitatively mirrored those of Experiment 1, showing that only Controls tend to show more yes-response advantage in metacognitive performance with HSF relative to LSF stimuli, albeit non-significant. See the main text for the statistical results. Error bars indicate standard error of the mean. HSF, High spatial-frequency, LSF, Low spatial-frequency.

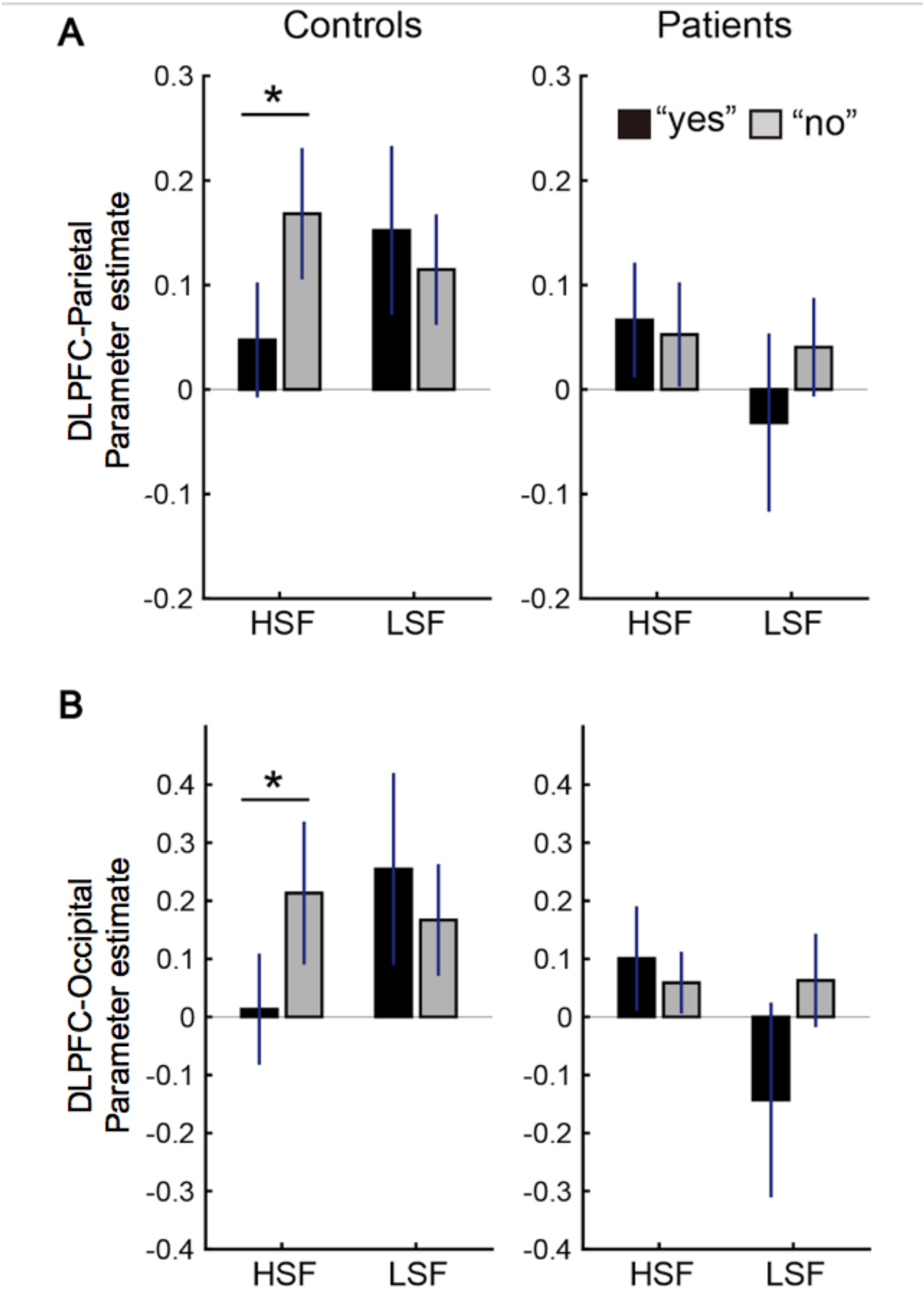


**Supplementary Figure 6.** The results of a general form of context-dependent psychophysiological interaction analysis (gPPI) with DLPFC as a seed ROI. As shown in Figure 3, the functional connectivity between DLPFC and bilateral parietal areas as well as occipital areas were significantly modulated as a function of interaction between Group, Spatial-frequency, and Response-type ($p < 0.01$, corrected with cluster-size thresholding). **A.** Parameter estimates (Beta) for the gPPI analysis within the parietal area for each Group, Spatial-frequency level, and Response-type. **B.** Parameter estimates for the gPPI analysis within the occipital area for each condition. For both parietal and occipital clusters, Controls showed a significant difference in parameter estimate between yes- and no-response trials with HSF stimuli but not with LSF stimuli. Patients showed no significant difference in parameter estimates across conditions. a: anterior, r: right, * $p < .05$. *Related to Figure 3.*

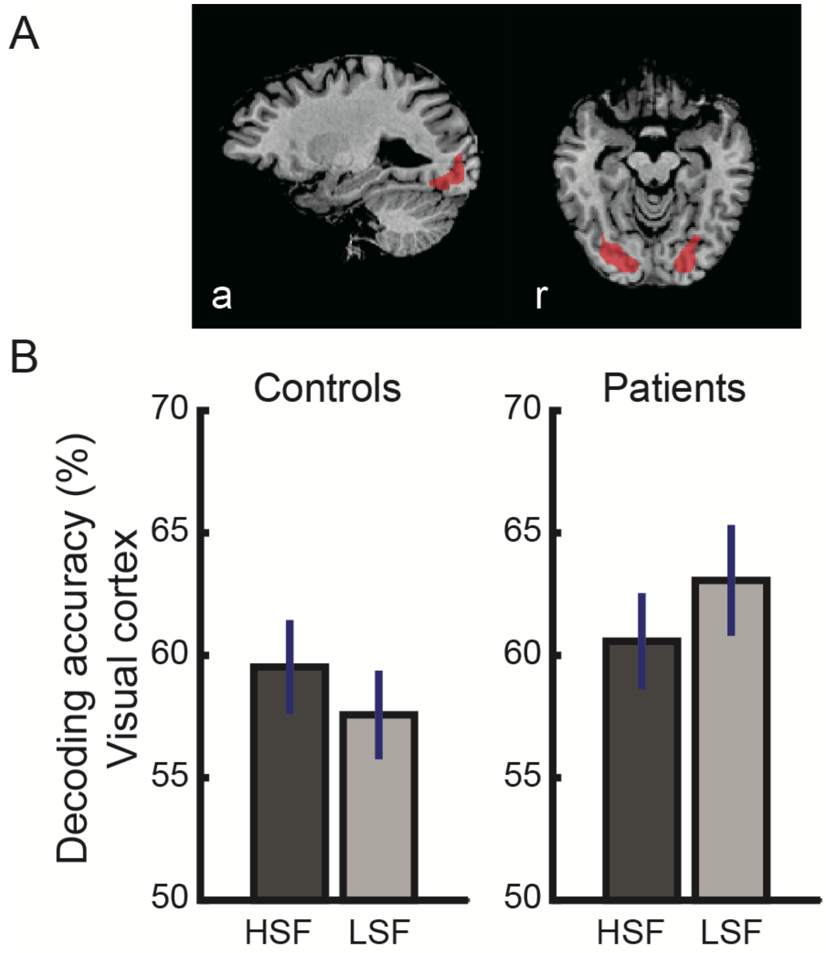


**Supplementary Figure 7.** Control ROIs in visual cortex and its decoding results. **A.** Control ROIs functionally defined from a group GLM. ROIs include the voxels that showed significantly larger activity during the stimulus period relative to fixation in a group GLM ($p < .01$, Bonferroni corrected). **B.** Decoding accuracy of confidence rating (high versus low) within Control ROIs. There were no significant main effects of Group and Spatial-frequency, and their interaction (see Results for details). Error bars indicate standard error of the mean. a: anterior, r: right, HSF: High spatial-frequency, LSF: Low spatial-frequency. Related to Figure 4.

### Pilot Experiment

We conducted Pilot Experiment to establish the procedure for Experiment 1 and 2.

#### Methods and Materials

*Participants*

Fifteen students from Columbia University were enrolled. They had either normal or corrected-to-normal vision, and were free from clinical conditions. All participants gave written informed consent prior to their participation and received $10/h. The study was approved by the Columbia University's Committee for the Protection of Human Subjects. The estimated age range is from 18 to 23. Further details of their demographic information are missing due to research site relocations.

*Procedure*

The task consisted of a detection task block (720 trials, performed in eight subblocks of 90 trials each), which followed two practice blocks (28 trials each) and one calibration block (120 trials). On each trial of the main blocks (Figure 1A), a fixation cross was presented on the center of the screen for 1 s, replaced by a background patch (4° in visual angle) containing dynamic white noise refreshed at 60 Hz (similar to TV static) presented for 1 s. On half of the trials, a horizontal grating (2.6 cycle per degree, cpd) briefly emerged within a background noise patch for 250 ms. On the other half of trials, a background patch remained for another 250 ms without a grating. For both trial types, the noise patch alone continued to be presented for a final 250 ms. Subsequently, participants were asked to indicate whether there was a grating (yes-response) or not (no-response) by pressing either "1" or "2" key. The key-assignment was counterbalanced across participants and was fixed across the blocks for each participant. They were further asked to rate their confidence in perceptual response on a four-point scale, by pressing one of the aligned 7-8-9-0 keys on a keyboard. The assignment for confidence rating was fixed across participants to minimize task load (i.e., assigning higher confidence to smaller number, rather than to larger number, would unwantedly increase task load). For both perceptual response and confidence rating, response-to-key assignments were presented on the lower screen as reminders, and participants made responses within 5 s. The next trial began after participants responded or 5 s elapsed from the onset of reminder for confidence rating. The trial sequence of the calibration blocks was identical to that of the main blocks, except that a horizontal grating appeared on all the trials.

During the calibration block, Michelson contrast of grating was titrated with QUEST threshold estimation procedure [15]. Three independent sequences of 40 trials were interleaved during the calibration block to yield three estimates of contrast level to achieve hit rate of 50%. The median of three estimates was multiplied by 0.7, 1.0, 1.3 to serve as three levels of difficulty (high/mid/low) in the detection task block. There were equal numbers of trials with high, mid, and low task difficulty levels. The order of trials was randomized within each block.

The background of the screen remained gray throughout the blocks, and the viewing distance was fixed at 60 cm. The stimuli were generated and presented with the Psychophysics Toolbox (Watson and Pelli, 1983) in MATLAB (Mathworks) implemented in iMac (21.5 inch).

*Analyses*

Detection sensitivity was calculated as d' for each of the three levels of task difficulty with standard signal detection theory (SDT) methods (Macmillan and Creelman, 1991). Metacognitive sensitivity for yes- and no-responses was separately calculated as response-specific meta-d' (rs-meta-d'; http://www.columbia.edu/~bsm2105/type2sdt/fit_rs_meta_d_MLE.m) (Maniscalco and Lau, 2014), separately for each of the three levels of task difficulty. We estimated rs-meta-d' with the assumption of equal variance between signal (i.e., target presence) and noise (i.e., absence) (i.e., z-transformed receiver operating characteristic (zROC) slope, s = 1) (see related technical issues in properly correcting for potential unequal variance in detection tasks (Maniscalco and Lau, 2014)). The ratio of meta-d' to d' (meta-d'/d') was then calculated for each difficulty level, to quantify metacognitive efficiency i.e., how much sensory signal remained available for metacognitive judgement given how much was originally accessible for perceptual judgement (Fleming and Lau, 2014; Maniscalco and Lau, 2014). Across-participant means for d', meta-d', and meta-d'/d' were analyzed with Analysis of Variance (ANOVA) in SPSS version 25 (IBM). One caveat of the ANOVAs here was that the same set of target absent trials were resampled to estimate d' and meta-d' across different task difficulty levels. However, we considered this as only a minor issue as it did not bias certain experimental conditions over others. For the subsequent analyses, an ANOVA with two within-participant factors of Response-type (yes/no) and Task-difficulty (high/mid/low) were conducted.

**Results and discussion**

Means of response-specific meta-d'/d', response-specific meta-d', and detection d' are shown in Supplementary Figure 1 for each level of task difficulty. As shown in Supplementary Figure 1A, meta-d'/d' was generally higher for yes-responses than no-responses, although the main effect of Response-type was not significant ($F(1, 14) = 3.71$, $p = .075$). This yes-response advantage (i.e., a main effect of response-type) significantly interacted with Task-difficulty ($F(2, 13) = 4.27$, $p =.038$, partial $\eta 2 = .396$). This interaction was due to yes-response advantage being significant only with low difficulty level (i.e., highest contrast) ($p = .008$, Bonferroni-corrected) but not with mid and high difficulty levels ($p = .138$, $p = .217$, respectively).

As expected, detection d' (Supplementary Figure 1B) monotonically increased as the task difficulty level decreased (a main effect of Task-difficulty: $F(2,13) = 162.32$, $p < .001$, partial $\eta 2 = .961$). That is, low difficulty level yielded significantly higher detection d’ than mid difficulty level ($p < .001$), and mid difficulty level yielded significantly higher detection d’ than high difficulty level ($p < .001$).

The analysis of meta-d' (Supplementary Figure 1C) was qualitatively similar to that of meta-d'/d', revealing a significant main effect of Response-type ($F(1, 14) = 4.76$, $p = .047$, partial $\eta 2 = .254$) which significantly interacted with Task-difficulty ($F(1, 13) = 6.72$, $p = .01$, partial $\eta 2 = .508$). Similar to the aforementioned results of meta-d'/d', this interaction was due to the fact that yes-response advantage was significant only with low task difficulty level ($p = .006$, Bonferroni-corrected) but not with mid or high task difficulty levels ($p = 225$, $p = .276$, respectively).

Taken together, the results suggest that yes-response advantage in metacognition is robust enough to be qualitatively retained across different levels of task difficulty (i.e., stimulus contrast), although it was only significantly observed with the easiest task difficulty (i.e., highest contrast), potentially due to lowered metacognitive performance even for yes-response with increased task difficulty. Such a yes-response advantage is in line with previous studies (Fleming and Dolan, 2010; Kanai et al., 2010), suggesting that response-specific meta-d' used here has sensitivity to capture the difference in metacognitive sensitivity between yes- and no-responses.

### ***Additional information on Experiment 1***

*Participant*

Patients were recruited through word-of-mouth via the Department of Psychiatry, Kyoto University Graduate School of Medicine. Healthy individuals were recruited through word-of-mouth via Center for Information and Neural Networks (CiNet), National Institute of Communications Technology (NICT). The protocol was approved by the ethical committee of Kyoto University as well as by that of NICT. All participants gave written consent form prior to participation. They had normal or corrected-to-normal visual acuity. Participants received around $40 (in Japanese yen) for their participation unless not eligible (e.g., participating in paid hours or on exclusive scholarship).

The patients included in analyses were medicated with antipsychotics (atypical N = 15; typical + atypical N = 2). The patients and control subjects did not statistically differ in their age ($t(33) = .19$, $p = .850$) and gender ratio ($\chi 2 = .69$, $p = .407$). However, consistent with previous studies and the nature of schizophrenia (Calderone et al., 2013), patients showed significantly lower estimated IQ ($M = 108.2 \pm 7.8$) than control subjects ($M = 112.5 \pm 5.0$) ($t(33) = 4.08$, $p < .001$), when measured with a standardized Japanese Adult Reading Test (JART-25) to estimate full scale intelligence quotient (FSIQ) (Matsuoka and Kim, 2007; Matsuoka et al., 2006).

*Procedure details*

Participants performed a pair of tasks to detect a grating in HSF and that in LSF (2.6 cycle per degree, cpd; 0.4 cpd, respectively), in a counterbalanced order. Detection tasks with HSF and LSF grating targets were conducted in two separate sessions scheduled on two separate days. Each task consisted of a detection task block (360 trials, performed in six subblocks of 60 trials each), which followed two practice blocks (10 trials each) and one calibration block (80 trials). On each trial of the main blocks (Figure 1A), a fixation cross was presented on the center of the screen for 1 s, replaced by a background patch (4° in visual angle) containing dynamic white noise refreshed at 60 Hz (similar to TV static) presented for 1 s. On half of the trials, a horizontal grating briefly emerged within a background noise patch for 250 ms. Gratings were embedded in a noise patch (diameter = 3.9° in visual angle). These two levels of SF were expected to evoke differential neural activity between patients with schizophrenia and control subjects (Martinez et al., 2008). On the other half of trials, a background patch remained for another 250 ms without a grating. For both trial types, the noise patch alone continued to be presented for a final 250 ms. Subsequently, participants were asked to indicate whether there was a grating (yes-response) or not (no-response) by pressing either “1” or “2” key. The key-assignment was counterbalanced across participants and was fixed across the blocks for each participant. They were further asked to rate their confidence in perceptual response on a four-point scale, by pressing one of the aligned 1-2-3-4 keys on a keyboard to enable the participants to make both perceptual and metacognitive responses with only one hand, avoiding potential difficulty in coordinating two hands among patients. The assignment for confidence rating was fixed across participants to minimize task load (i.e., assigning higher confidence to smaller number, rather than to larger number, would unwantedly increase task load). For both perceptual response and confidence rating, response-to-key assignments were presented on the lower screen as reminders, and participants made a response within 5 s. The next trial began after participants responded or 5 s elapsed from the onset of reminder for confidence rating. The trial sequence of the calibration blocks was identical to that of the main blocks, except that a horizontal grating appeared on all the trials.

During the calibration block, Michelson contrast of grating was titrated with QUEST threshold estimation procedure (Watson and Pelli, 1983). Two sets of 40 trials were randomly interleaved to estimate the two levels of contrast to achieve hit rate of 55% and 65%. Calibration was conducted separately for each SF level. Accuracy in the initial 10 trials of each set was disregarded, so as to discern irrelevant errors before accustomed to the task procedure. The two contrast levels were alternated in the subsequent six main blocks in order to avoid over-habituation to the stimuli with fixed contrast. To compensate for the potential fluctuation in performance due to fatigue or learning during the main blocks, the contrast levels were adjusted by a small increment (.015) to slightly increase or decrease the contrast when the mean response accuracy was too low (< 45%) or too high (> 65%) in a given main block. We here allowed a liberal range of performance (i.e., 45% to 65%) to be kept without adjustment in order to avoid overcorrection of the contrast levels between the main blocks. The contrast level of grating was fixed within a given main block. The phase of grating was randomized across trials. Confidence rating was omitted during the calibration block to minimize the overall task load, especially for the patients.

The background of the screen remained gray throughout the blocks, and the viewing distance was fixed at 55 cm. The order of trials was randomized within each block. The stimuli were generated and presented with the Psychophysics Toolbox (Brainard, 1997) in MATLAB (Mathworks) implemented in iMac (13 inch).

*Analyses*

The perceptual and metacognitive performance was estimated as d' and meta-d' as in Pilot Experiment, while collapsing the relatively small across-block variability in grating contrast. We conducted repeated measures ANOVAs with Response-type (yes/no) and Spatial-frequency (HSF/LSF) as within-subject factors and Group (Patients/Controls) as a between-subject factor. Unlike in Pilot Experiment, there was no violation of value dependence in Experiment 1, as there were different sets of target-absent trials for each condition (e.g., target spatial-frequency).

We estimated rs-meta-d' with the assumption of equal variance between signal (i.e., target presence) and noise (i.e., absence) (i.e., z-transformed receiver operating characteristic (zROC) slope, s = 1) (see related technical issues in properly correcting for potential unequal variance in detection tasks (Maniscalco and Lau, 2014)).

Metacognitive sensitivity for yes- and no-responses was separately calculated as response-specific meta-d' (rs-meta-d'; http://www.columbia.edu/~bsm2105/type2sdt/fit_rs_meta_d_MLE.m) (Maniscalco and Lau, 2014), separately each SF level. The ratio of meta-d' to d' (meta-d'/d') was then calculated for each difficulty level to quantify metacognitive efficiency (Fleming and Lau, 2014; Maniscalco and Lau, 2014). Meta-d’ corresponds to the value of d’ that would be expected to generate a subject’s measured level of metacognitive sensitivity, according to SDT. Thus, for a subject whose behavior matches SDT expectation, meta-d’ = d’, meaning that the subject’s measured metacognitive sensitivity (meta-d’) corresponds to the level of metacognitive sensitivity that would be expected to arise from their performance on the primary perceptual task (d’). For subjects with meta-d' < d', metacognitive sensitivity is lower than would be expected based on primary task performance. For such subjects, metacognitive sensitivity can be considered to be "inefficient" or "suboptimal," relative to SDT expectation.

Additional results

For the detection sensitivity d' (Supplementary Figure 3), there was a main effect of SF ($F(1, 33) = 4.33$, $p = .045$, partial $\eta 2 = .116$), which was due to generally lower d' with LSF relative to HSF stimuli. Note that this main effect alone cannot account for the aforementioned group difference in metacognitive performance because of at least two reasons. First, meta-d'/d' already takes the variability in d' into account when assessing metacognitive efficiency (Maniscalco and Lau, 2014). Second, if the difference in d' accounts for the results in meta-d'/d', then we would expect that the result of meta-d'/d' (i.e., yes-response advantage) would differ between HSF and LSF stimuli for Patients rather than for Controls. This is because, although the interaction was non-significant, it was Patients who primarily showed difference in d' between HSF and LSF stimuli (Supplementary Figure 3). However, the results of meta-d'/d' showed that it was Patients who showed equivalent yes-response advantage between SF levels.

The calibrated stimulus contrast tended to be higher for Patients relative to Controls (M = 9.79% ± s.d. 2.78, M = 8.27% ± 1.55, respectively), but did not statistically differ across Groups (a main effect; $F(1,33) = 4.05$, $p = .052$). There was no significant main effect of SF (a main effect; $F(1,33) = 2.43$, $p = .135$) or two-way interaction ($F(1, 33) = .18$, $p = .677$). The analysis of meta-d' (Supplementary Figure 3) alone revealed a qualitatively similar result as that of meta-d'/d'.

*Relationship between hallucination severity and metacognitive performance*

Among Patients in Experiment 1, there was further variability in their metacognitive performance as a function of their hallucination severity assessed with PANSS score. Specifically, the hallucination severity was generally related with weaker yes-response advantage in metacognition, although this correlation was significant only with LSF stimuli (r = -.516, p = .034) but not with HSF stimuli (r = -.367, p = .147). This result, however, should be treated cautiously as the hallucination severity was also significantly correlated with detection d' with LSF stimuli (r = .528, p = .029), although not with HSF stimuli (r = .289, p = .260). When d' was included as a covariate, the hallucination severity no longer correlated with the yes-response advantage in both HSF and LSF stimuli (r = -.265, p = .321; r = -.382, p = .144, respectively). Future studies may re-examine the replicability of the relationship between metacognitive performance and hallucination severity, with tighter stimulus calibration. The correlation between hallucination and d' is potentially due to that stimulus was calibrated solely based on hit rate rather than on d' (see *Procedure*). This may have resulted in less successful stimulus calibration among more severe hallucinators who potentially had lowered hit rate not because of perceptual difficulty but because of response bias, resulting in higher stimulus contrast. Indeed, patients with more severe hallucinations had higher stimulus contrast albeit with calibration (i.e., a positive correlation between the hallucination severity and stimulus contrast with LSF; r = .543, p = .024, although it did not reach significance with HSF; r = .402, p = .109).

### ***Additional information on Experiment 2***

*Participant recruitment*

Patients were recruited through word-of-mouth via the Department of Psychiatry, Kyoto University Graduate School of Medicine. Healthy individuals were recruited through word-of-mouth via Kyoto University as well as Center for Information and Neural Networks (CiNet), National Institute of Communications Technology (NICT). The protocol was approved by the ethical committee of Kyoto University as well as by that of NICT. All participants gave written consent form prior to participation. They had normal or corrected-to-normal visual acuity. Participants received around $75 (in Japanese yen) for their participation unless not eligible (e.g., participating in paid hours or on exclusive scholarship).

The patients included in the analyses were medicated with antipsychotics (atypical, N = 12, typical + atypical, N = 3). Those patients and controls did not statistically differ in their age ($t(30)=1.35$, $p = .186$) and gender ratio ($\chi2 = .74$, $p = .388$). As expected, the IQ estimated with JART [23, 24] was significantly higher for controls (M = 110.5 ± 5.7) than for patients (M = 98.9 ± 9.0) ($t(30) = 4.42$, $p < .001$). Nine patients and seven controls were also enrolled in Experiment 1, and the ratio of overlapped participation did not statistically differ between the groups ($\chi2 = 1.13$, $p = .288$).

*Procedure details*

First, the detection task with each stimulus frequency level (HSF or LSF) was divided into four runs of 24 trials. Second, the trial sequence was modified to be in synchrony with TRs, especially the onset of confidence rating period which is of primary interest here. Specifically, each trial started with a counterbalanced fixation period of 3 or 5s, followed by a stimulus period of 2.6s (1s noise + 1.5s grating or noise + 0.1s noise). The stimulus period was followed by 2.4s response period to indicate the presence of grating (yes or no), which was followed by 4s respond period to indicate the confidence level (1 to 4). The stimulus was enlarged to a diameter of 9° (in visual angle) to ensure sufficient neural activity.

*fMRI measurements*

Participants were scanned in a Siemens Trio 3 Tesla MRI scanner (Erlangen, Germany) with 32 channel head coil installed in Human Brain Research Center, Kyoto University Graduate School of Medicine (Kyoto, Japan) or Center for Information and Neural Networks (CiNet) (Osaka, Japan). All Patients and nine Controls were scanned in the former scanner. The remaining eight

Controls were scanned in the latter scanner to avoid conflict with other medical use of the former scanner. The fMRI parameters were identical between the locations: For functional echo-planar image (EPI) acquisition; TR = 2000 ms, TE = 30 ms, flip angle = 70 deg, FOV = 192 mm, Slices = 50, voxel size: 3 x 3 x 3 mm, Multi-band accel. factor = 2. An anatomical T1-weighted image was acquired with MPRAGE sequence; TR = 1900 ms, TE = 2.48 ms, flip angle = 9 deg, voxel size: 1 x 1 x 1 mm.

*Analyses*

The data were corrected for their slice timing and 3D head motion, and went through temporal high pass filtering. The data were not spatially smoothed for the multivoxel decoding purpose, but were smoothed with a kernel of 6 mm for the purpose of group GLM (see *ROI definition in Main text*). Individual T1 images and EPI were normalized to Talairach coordinate.

*Behavioral results*

Overall, the result of Meta-d'/d' qualitatively mirrored that of Experiment 1. Yet, the result in Experiment 2 was noisier (i.e., larger variability), which was well expected due to fewer trial numbers, tighter response time constraint, and being in a physically constrained fMRI (Supplementary Figure 5). That is, Meta-d'/d' was numerically larger for yes than no-responses (i.e., yes-response advantage), although this difference did not reach significance (a main effect of Response-type, $F(1, 30) = 3.39$, $p = .075$). Only Controls showed numerically larger Meta-d'/d' for yes- than no-responses (i.e., yes-response advantage) with HSF (Δ Meta-d'/d': M = 0.908 ± s.e. 0.489, $p = .175$) but not with LSF (M = -0.009 ± s.e. 0.212, $p = .975$). Meanwhile, Patients showed similar yes-response advantage with HSF (M = 0.725 ± 0.853, $p = .306$) and LSF (M = 0.460 ± 0.367, $p = .136$). There was no significant interaction between SF, Response-type, and Group ($F(1,30) = .48$, $p = .493$).

The detection sensitivity d' did not differ as a function of SF ($F(1, 30) = .02$, $p = .899$), that of Group ($F(1, 30) = 1.39$, $p = .247$), and their interaction ($F(1, 30) = .13$, $p = .721$). This result suggests that calibration of stimulus contrast was comparable across all experimental conditions. The mean contrast levels of Controls and Patients were 5.4% ± s.e. 0.42 and 4.78% ± 0.46, respectively. There was neither a significant main effect of Group ($F(1,30) = .98$, $p = .330$) nor that of SF ($F(1, 30) = 2.80$, $p = .105$). The interaction between Group and SF was also non-significant ($F(1, 30) = 1.09$, $p = .305$).

*ROI definition*

The clusters in the left and right hemispheres were combined to form the DLPFC ROI for subsequent analyses. Note that bias in voxel selection for any particular experimental condition (e.g., HSF) or group was minimized, as all the trials including all the conditions from all participants were included in a GLM. To further ensure this, we ran an analysis of covariance (ANCOVA) examining the effects of Group, SF, Response-type in BrainVoyager to show that no voxel survived the cluster-threshold enhancement (Forman et al., 1995; Goebel et al., 2006) at liberal criteria ($p < .01$) with a mask of DLPFC ROI. This was consistently the case when examining a main effect of Group, SF, Response-type, and the interactions among any combinations of the variables.

For the DLPFC ROI, the center of gravity in Talairach space was [X = -34.28 (± SD 2.45), Y = 35.33 (± 3.37), Z = 32.43 (± 3.81)] for the left hemisphere cluster and [X = 37.67 (± 3.52), Y = 30.44 (± 3.43), Z = 31.61 (± 3.35)] for the right hemisphere cluster. The number of voxels was 994 and 1425 in the left and right hemispheres, respectively. For the control ROI within visual cortex, the center of gravity in Talairach space was [X = -22.21 (± SD 3.88), Y = -81.09 (± 5.47), Z = -10.29 (± 5.03)] for the left hemisphere and [X = 22.40 (± 5.96), Y = -81.35 (± 5.56), Z = -8.27 (± 2.78)] for the right hemisphere, which partially overlaps with the anatomical landmarks of V4 (Witthoft et al., 2014). The number of voxels was 2833 and 2339 in the left and right hemispheres, respectively.

*Generalized form of context-dependent psychophysiological interaction analysis (gPPI)*

We aimed to examine whether functional connectivity between DLPFC and some other brain areas were modulated as a function of Spatial-frequency, Response-type, and Group. With this aim, we first conducted a whole-brain gPPI analysis (McLaren et al., 2012) with DLPFC as a seed ROI to quantify its functional integration with other brain areas as a function of Spatial-frequency and Response-type at the participant level. The parameter estimates (Beta values) for PPI terms were then analyzed with a group-level ANOVA to examine whether the functional connectivity between DLPFC and elsewhere in the brain may be modulated as a function of Spatial-frequency, Response-type, and Group. The whole-brain gPPI model included regressors for each of the four conditions (i.e., yes- and no-responses with HSF and LSF stimuli) convolved with the Boynton HRF (Boynton et al., 1996), a regressor for the z-normalized time course of the seed DLPFC ROI, regressors for PPI terms (i.e., condition regressor x seed time course), as well as six nuisance regressors of 3D head motion including 3 translation directions and 3 rotation axes.

The whole-brain gPPI analysis revealed the bilateral clusters in parietal cortex and visual cortex located posterior of BA 7 and ventral of BA18 (MRIcron, Brodmann 48 Area Atlas Template), respectively. The centers of gravity for the left and right clusters in parietal cortex were [X = -21.34 (± 4.76), Y = -68.37 (± 4.18), Z = 38.32 (± 3.40)] and [X = 10.50 (± 2.28), Y = -61.86 (± 3.13), Z = 36.32 (± 2.41)], respectively. The centers of gravity for the left and right clusters in visual cortex were [X = 17.13 (± 4.62), Y = -72.81 (± 2.71), Z = -16.73 (± 2.92)] and [X = -24.0 (± 5.21), Y = -77.51 (± 3.22), Z = -19.52 (± 1.93)], respectively. There were also bilateral significant clusters in motor cortex overlapping BA 6 (Figure 3), which may reflect mere motor-related consequences of the metacognition-level effect (e.g., reduced motor fluency in responding with difficult metacognitive judgement).

*Decoding analysis*

To binary decode the trial-by-trial confidence level (high or low) from the multivoxel activation pattern in DLPFC during the confidence rating period, we built a decoder with sparse logistic regression (SLR) which automatically selects relevant features (i.e., voxels) (Yamashita et al., 2008). SLR uses a linear discriminant function (LDF) to separate two classes, S1 and S2 (high and low confidence, respectively), based on the weighted sum of the value for each feature (i.e., voxel) (see (Cortese et al., 2016; Yamashita et al., 2008) for details). We used SLR instead of other classifying algorithms such as support vector machines as SLR has been demonstrated to be advantageous when the size of features (voxels) are much larger than that of samples (trials) (Yamashita et al., 2008). Specifically, to handle a sample size smaller than the size of features (voxels), irrelevant voxels were automatically pruned (see *Details on SLR* and Yamashita et al. (Yamashita et al., 2008) for more detailed descriptions).

The multivoxel activation pattern per trial was extracted by averaging the activation patterns in the DLPFC ROI across the 6s time window (i.e., 3 TRs) covering the confidence rating period from its onset while considering a haemodynamic delay of 3TR. The time course data were z-transformed prior to averaging. We averaged the patterns across the 6s time window to stabilize the patterns, as in previous studies (Amano et al., 2016; Cortese et al., 2016; Koizumi et al., 2016).

The data from the correct trials with confidence rating of 1 or 4 were sent to the decoding analysis to represent a class of low and high confidence, respectively. We here only included the correct trials as in a previous study (Cortese et al., 2016) in order to avoid building a decoder to discern correct versus incorrect trials, which generally coincide with higher and lower confidence trials, respectively. When the trial number of either class was smaller than the other, the shortage was covered by additionally including the randomly sampled trials with confidence rating of 2 or 3. As this random sampling could jitter the results of the decoding analysis, the process was repeated ten times, each time with renewed random sampling. The results were then averaged across the repetitions per participant. Within each repetition of the decoding analysis, the SLR-classification was optimized through an iterative approach. Specifically, a decoder was built with 10 iterations, where only the previously unselected features were used in the next iteration. Here, the samples were divided into a training set and a test set, where a training set was used to build a decoder and a test set was used to estimate the accuracy of the built decoder while avoiding overfitting. The optimal number of SLRs (iterations) was then used to estimate the decoding accuracy. The data from one Control participant was excluded for having too few trials with confidence rating of both 1 and 4. We were unable to run an analysis separately for yes- and no-response trials due to insufficient trial numbers in each class of confidence level.

The decoding accuracy was analyzed with a Group x Spatial-frequency ANOVA. Meta-d'/d' for each Spatial-frequency level (averaged across response-types) was included as covariates of uninterest, in order to exclude the possibility that variability in decoding accuracy of confidence level simply reflected that in metacognitive performance.

*Details on SLR*

SLR uses a linear discriminant function (LDF) to discriminate two classes, $S_1$ and $S_2$ (High and Low confidence, respectively), while using the weighted sum of the value from each of the features (voxels),

$$f(x;\theta) = \sum_{d=1}^{D} \theta_d x_d + \theta_0. \quad (1)$$

Where $x = (x_1, \ldots, x_D)^t \in R^D$ is the input vector in each feature (i.e., BOLD signal in a voxel) in D dimensional space. And $\theta = (\theta_0, \theta_1, \ldots, \theta_D)^t$ is the weight vector which includes a bias term ($\theta_0$).

The boundary dividing the two classes is the hyperplane where $f(x;\theta) = 0$. Logistic regression outputs the likelihood of class $S_1$ (i.e., High confidence) given an input feature with the logistic function,

$$p = \frac{1}{1+exp(-f(x;\theta))} \equiv P(S_1|x)\,. \tag{2}$$

Here, $p$ would equal to 0.5 when $f(x;\theta) = 0$ (i.e., the boundary), where $p$ approaches 0 or 1 when $f(x;\theta)$ approaches to plus or minus infinity, respectively. With our data set, logistic regression was not directly applicable because it had a smaller number of samples relative to that of features. Thus, we reduced the dimensionality by automatically pruning out irrelevant voxels. For more detailed descriptions of this reduction procedure, see Yamashita *et al (Yamashita et al., 2008).*
